# Conservation of extended sequence and structure in the branchpoint-to-3’ splice site region upstream of neural microexons

**DOI:** 10.64898/2026.01.16.699960

**Authors:** Alexandra Randazza, Kathryn E. Howe, John R. McCoy, Abigail Hatfield, Tara Doucet-O’Hare, Lela Lackey

## Abstract

Microexons are short exons that are highly conserved in vertebrates and are essential for neurodevelopment. Their small size poses a challenge for regulatory protein binding and exon-definition splice site recognition, which typically relies on standard length exons. Here, we determine the sequence and RNA structural features of neural microexons in humans and in the chick developmental model organism. We demonstrate that a subset of neural microexons undergoes dynamic, stage-specific regulation during chick embryonic brain development that correlates with expression of known microexon regulators, *SRRM4* and *NOVA1*. Using experimental RNA structure-probing on a subset of neural microexons, we show that shared RNA secondary structures between orthologous human and chicken microexon precursor mRNAs primarily occur in regions of high sequence conservation. We find that both human and chicken neural microexons have extended functional distance between the branchpoint and the 3’ splice. Structurally, branchpoint-to-splice site regions are unusually accessible and relatively unpaired compared to other exon classes. Our data suggest that microexon splicing relies on structural accessibility of the branch-point-to-splice site region, which may influence accessibility for SRRM4 binding and alleviate steric constraints for spliceosome assembly.

**Graphical Abstract:** 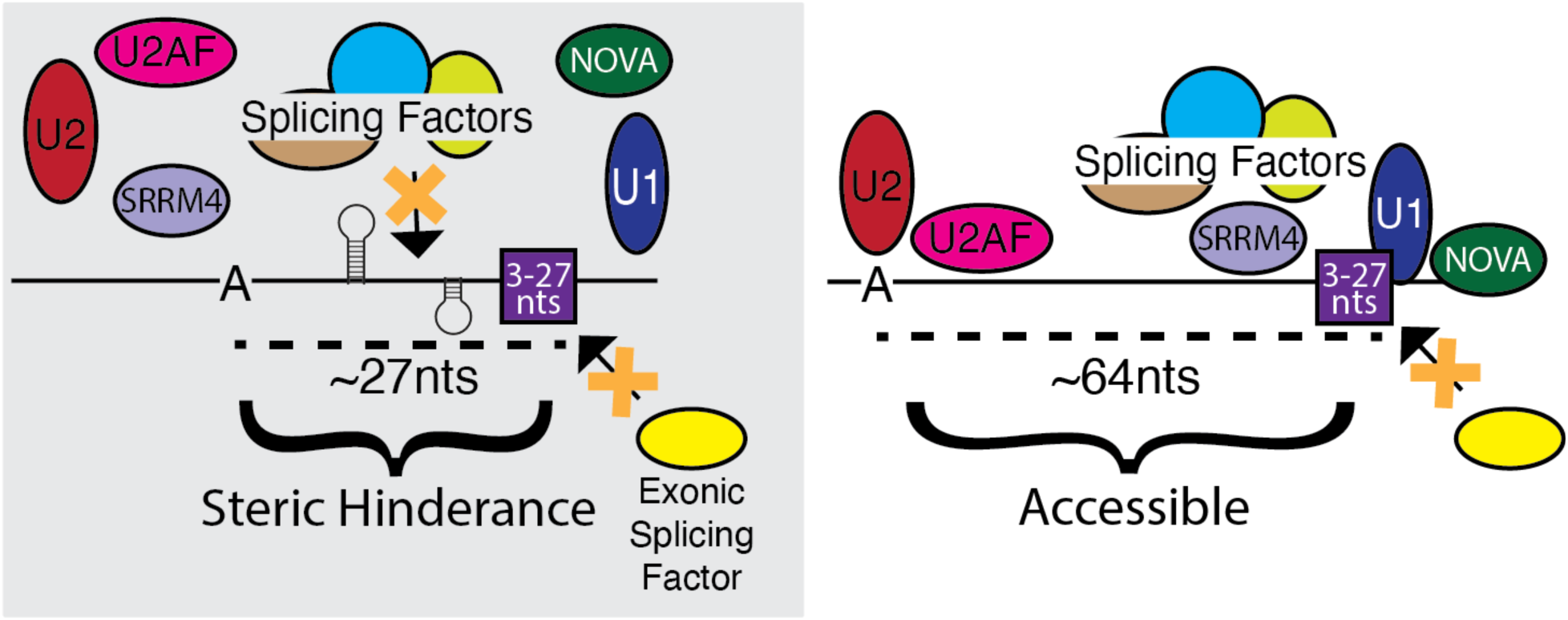

## Introduction

The brain exhibits some of the highest levels of alternative splicing in the human body, producing a diverse array of RNA isoforms, including transcripts containing microexons (1). Microexons are short exons (3-27 nucleotides (nts)) that are highly conserved in vertebrates (2–4). Microexons are usually part of the coding sequence and retain open reading frames, frequently resulting in small amino acid insertions in proteins. These insertions can alter protein binding affinities and partners (3,5). Specific microexons have been linked to neurological disorders and development (5), including a microexon in cytoplasmic polyadenylation element binding protein 4 (*CPEB4*) that broadly regulates translational efficiency and is associated with autism (6–8), and a microexon in roundabout guidance receptor 1 (*ROBO1*) that influences neural migration during brain development (9). Changes in microexon splicing can be caused by misregulation of the splicing factor serine/arginine repetitive matrix 4 (SRRM4) protein and have been associated with a variety of neurological disorders, such as autism and schizophrenia (3,10,11).

Splicing depends on both sequence and structural elements to guide splice site selection and promote efficiency (12–14). For example, hairpin formation at the 5’ splice site of the microtubule associated protein tau (*MAPT*) precursor RNA can inhibit spliceosomal recognition and is essential for appropriate splice site recognition and balance of mature mRNA isoforms (15). However, RNA structure does not simply block protein binding, studies of early spliceosome assembly on hairpin containing precursor RNA reveal that hairpins can also promote splicing (16). Our previous work studying the secondary structure of precursor RNAs found a key role for RNA flexibility in splicing fidelity, with mis-spliced RNAs being much more flexible than robustly spliced RNAs (17). However, the structure of most precursor RNAs, including precursor RNAs containing microexons, remains unknown. Furthermore, how structural and sequence elements interact to influence splice site selection is unclear.

In vertebrates the exon-definition model of splice site selection depends on the fact that exons are 120-150 nts in length, which makes them more consistent and significantly shorter than most vertebrate introns at 1,100-1,700 nts in length (18,19). In the exon-definition model, three splice sites are identified, starting with the original 5’ splice site and the downstream 5’ splice site. The downstream 5’ splice site then assists with identification of the upstream 3’ splice site. The spliceosome then remodels into an intron-spanning complex for splicing with the original 5’ splice site and the newly identified 3’ splice site (16,20,21). Exon definition is assisted by members of the family of serine-arginine rich RNA binding proteins (RBPs), including serine and arginine rich splicing factor 1 (SRSF1), that can bind exon enhancer elements or be recruited by the U1 small nuclear ribonucleoprotein (snRNP) component of the spliceosome (22). This mechanism requires significant space for assembly of splicing factors at the 5’ splice site and subsequent recruitment of splicing factors at the upstream 3’ splice site. One way to circumvent traditional exon-definition is through binding of regulatory factors that recruit the spliceosome. The microexon splicing factor SRRM4 is a serine-arginine rich RBP that regulates multiple microexons through a UGC binding motif and promotes microexon splice site recognition (23). Another possibility is that RNA structure can alter the distance between splice sites or regulatory elements (24–26), potentially influencing intron or exon definition splicing. Given their small size, microexons fall outside the bounds of conventional exon-definition splicing and may require structural and/or RBP co-factors to overcome these size constraints (27).

Here we analyze the properties of neural-specific microexons in humans and in chickens. Developmental milestones in the chicken embryo are well characterized, providing a unique opportunity to explore microexon conservation in a developmental context (28). We show that chicken microexons have dynamic temporal regulation during embryonic brain development that correlates with expression of known microexon regulators, neuro-oncological ventral antigen 1 *(NOVA1)* and *SRRM4*. A significant subset of these regulated microexons contain UGC binding motifs for SRRM4. We find that both human and chicken neural microexons have extended intronic polypyrimidine tracts that push the predicted branch-point farther into the intron. In addition, both computational and experimental structure models indicate that microexons have less base-pairing between the branch-point and 3’ splice site than standard-size exons. Our results suggest that, despite their small size, microexons are robustly spliced due to their extended branchpoint-to-splice site distance, which may relieve steric hinderance for co-factors like SRRM4 and spliceosomal proteins.

## Materials and Methods

### Chicken brain and heart sample collection

Tetra-H chicken (*Gallus gallus domesticus*) eggs were incubated at ∼37.5 °C, monitored by thermometer, for 42 hours, 56 hours, 5, 7 and 9 days at a ∼50% humidity to reach Hamburger-Hamilton (HH) stages 11, 15, 27, 31 and 35, respectively (29). Initially, fertilized eggs obtained from the Clemson Morgan Poultry Center (Clemson, SC) were stored at 15 °C for up to one week, and before incubation eggs were left at room temperature for 24 hours. Eggs were incubated in a Georgia Quail Farm (GQF) “Digital Sportsman” Model 1502 Incubator (Georgia Quail Farm, Cat. No. 1502) in humid conditions and were rotated mechanically every hour to facilitate embryo development. To collect embryos at HH stages 11 and 15, eggs were opened and the contents transferred to a Petri dish. Embryos were isolated from the yolk using a cloverleaf filter paper technique, and developmental stage was confirmed based on established Hamburger–Hamilton morphological criteria prior to tissue processing (29,30). The filter paper and embryo were transferred to a Petri dish with isotonic saline. The vitelline membrane was surgically removed to fully expose the embryo, and the heart was isolated and excised by sharp transection at its superior inflow and inferior outflow connections. The brain was subsequently dissected, and neural tissue encompassing the hindbrain and all more rostral regions were collected. After excision, both the brain and heart tissues were immediately transferred to TRIzol reagent (Invitrogen, Thermo Fisher Scientific), mechanically dissociated with pipetting, and flash frozen on dry ice. Embryos at HH stage 27 and later were harvested by opening the eggshell and transferring the embryo to a Petri dish containing isotonic saline. The vitelline membrane was carefully removed to expose the embryo. The heart was isolated and excised via a supra-aortic transection and immediately submerged in 500ul TRIzol reagent. For brain tissue collection, the ocular structures were removed along with surrounding non-neural tissues to fully expose the brain. Following gross dissection and cleaning of the neural tissue with saline, the brain was transferred to 500ul TRIzol reagent, mechanically dissociated using sterile scissors, and flash frozen. All tissue samples were stored at −80°C until RNA extraction.

### RNA extraction and library preparation

The samples stored in TRIzol were homogenized with 2.4mm metal bulk beads (Fisher Scientific) at 5 m/s for 60 seconds, repeating until tissue was fully homogenized. An additional 500ul TRIzol was added, and RNA extraction was performed following the manufacturer’s suggested protocol. After RNA isolation, samples were treated with DNase (Turbo DNA Free Kit, Invitrogen) according to the manufacturer’s instructions. Following standard ribosomal RNA depletion (Qiagen QIAseq FastSelect -rRNA HMR Kit or NEBNext rRNA Depletion Kit v2), cDNA was synthesized using the SuperScript™ IV First-Strand Synthesis System (Invitrogen, ThermoFisher Scientific) and Second Strand cDNA Synthesis Kit (Invitrogen, ThermoFisher Scientific) both according to the standard protocol. Libraries were prepared using the Nextera DNA Library Preparation Kit (Illumina). The paired-end libraries were sequenced at 2 x 150 on the Illumina NovaSeq. Samples were collected in triplicate per sex and per tissue.

### RNA sequencing analysis

Read quality was checked with FastQC v0.11.9 (https://github.com/s-andrews/FastQC). Reads were then mapped to the galGal6 genome assembly with STAR v2.7.10a (31) (Figure_1/star.sh) followed by filtering for uniquely mapped reads with Samtools v1.10 (32) (Figure_1/mapq_filt.sh). Aligned files were sorted and indexed with Samtools and gene expression was quantified with featureCounts from Subread v1.6.4 (33) followed by analysis with edgeR (34) (Figure_1/edgeR_analysis_plots.R). Splicing analysis was performed with rMATs v4.1.1 (35) (Figure_1/rmats_psi_heatmap.R) comparing splicing patterns between male and female derived tissues and between consecutive developmental stages for each tissue. Skipped exon events <= 27nts in length were included for further analysis with a special focus on the selected 13 microexons. RNA sequencing reads from induced pluripotent stem cell (iPSC)-derived neuronal progenitor cells (NPC) and neurons (GSE196144, 201B7 iPSC-derived NPC *n* = 3, 201B7 iPSC-derived neuron *n* = 3) were downloaded from the Sequence Read Archive (36) and processed using the same analysis pipeline after aligning to the Hg38 genome.

### Splice site selection

To develop lists of neural microexons we used VastDB tables “EVENT INFORMATION” and “MAIN PSI TABLE” datasets for human (hg38), chicken (galGal4), and mouse (mm10) (37). For each species, the VastDB events were filtered by size with microexons > 0 and <= 27 nts and midexons > 100 and <= 200 nts. Exon skipping events were selected. To ensure selection of mostly alternative exons, events with a percent spliced in (PSI) <= 5% and >= 95% in 4 or more samples were removed. Events with 10 or more reads in at least 50% of samples and 20 or more reads in at least 20% of samples were considered high confidence and used for further analysis. The samples from VastDB were divided into neural and non-neural categories. Welch’s t-test was used to identify exons that had significantly higher or lower PSI values in neural samples. After adjustment with Benjamini–Hochberg, events with p-values < 0.005 were classified as “neural” while those not meeting this threshold were classified as “non-neural” (event_filtering_criteria.R). UCSC LiftOver (38) was used to update the galGal4 coordinates to galGal6. Scripts for analyses and plots are available on GitHub (https://github.com/randazzal/extended_bp_2_3ss_scripts).

### Exon and intron characterization

Conservation was measured using phastCons scores (39) from UCSC Table Browser (assembly: hg38, group: Comparative Genomics, track: UCSC 100 Vertebrates, table: Cons 100 Verts (phastCons100way) (40) (Supplemental_Figures/PhastCons_A3ss_sample.R). Splice site strength was calculated with MaxEntScan (41) (Figure_3/splice_site_strength_human.R). Repeat content in upstream and downstream introns was generated with bedtools intersect (bedtools2 v2.29.2) (42) and repeat annotations for hg38 and galGal6 from UCSC Table Browser (track: RepeatMasker, table: rmsk) (43). Repeat content was also calculated with Tandem Repeat Finder (TRF) (44) using the recommended parameters (Supplemental_Figures /repetitive_sequence_plots.R). Branchpoints were identified using SVM-BPfinder (45) with “-s Hsap” and “-l 100” for both human and chicken due to the absence of a model trained for chicken sequences. The branchpoint with the highest “svm_scr” per intronic sequence was used for analysis (Figure_4/BPfinder_plots_human.R). UGC motifs were identified in fasta files and quantified using a custom python script (find_UGC.py). All analyses including sequence content (Figure_3/polypyrimidine_probabilities_human.R), and intron length (Supplemental_Figures /human_intron_length_plots.R) were conducted in R v4.1.2.

### Human and chicken sequence alignment

The VastDB table “EVENT CONSERVATION” was used to identify the chicken orthologs of selected human microexons and midexons. Sequences 200 nts upstream from the 3’ splice site and 200 nts downstream from the 5’ splice site were used for alignment. The global alignment score between orthologous flanking introns was calculated using PairwiseAligner from the Bio.Align package (https://biopython.org/docs/1.76/api/Bio.Align.html; Supplemental_Figures/alignment_score_seq_ident.ipynb) with default parameters. In addition, the unaligned sequence identity was calculated as “matches” divided by 200. Bootstrapping was performed in R v4.1.2.

### Conservation/covariation analysis

Multiple alignment files of the 450 nts in each direction from the 3’ splice site of a subset of neural microexons were downloaded from the multiz100way track (track: UCSC 100 vertebrates, table: Multiz Align (multiz100way) (46). The MAF files were converted to fasta files with Galaxy MAF to FASTA tool (47). Human (hg38), gorilla (gorGor3), Rhesus monkey (rheMac3), mouse (mm10), rat (rn6), cow (bosTau8), chicken (galGal4), western clawed frog (xenTro7), and zebrafish (danRer10) were kept and realigned using Clustal Omega v1.2.4 (48). Sequences on the human negative strand required an in-house python script to generate the reverse complement of all the sequences in the alignment (Figure_2/reverse_complement.ipynb). The ViennaRNA (49) package RNAalifold (50) was utilized to identify shared RNA structures between multiple species around the microexons using the following command: RNAalifold -p –maxBPspan 66 –sci –aln –color <alignment_fasta>.

### Chemical structure probing

Gene fragments for microexons and midexons of interest were designed and synthesized by Integrated DNA Technologies (IDT). Most human microexon and all midexon sequences contain 450 nucleotides (nts) flanking the 3’ splice site in both directions. Due to sequence constraints additional human microexons (*CPEB4, DCTN1, ASAP2, MEF2A, DOCK7*) and all chicken microexon sequences contain 200 nts flanking the 3’ splice site in both directions. A T7 promoter followed by a stabilizing hairpin sequence was on the 5’ end and a universal sequence was on the 3’ end of each gene fragment. The HiScribe T7 High Yield RNA synthesis kit (New England Biolabs, NEB) was used for *in vitro* RNA synthesis using the gene fragments acting as a template following the standard protocol. The synthesized RNA was treated with DNase I for 20 minutes at 37°C and cleaned up with RNA XP beads (Beckman Coulter). The RNA was incubated in folding buffer (400 mM bicine, 200 mM NaCl, 20 mM MgCL2), RNase inhibitor, and 5ul of 25 mM 5NIA or DMSO at 37°C for 5 minutes for SHAPE-MaP experiments (51). DMS-MaP experiments (52) used 5ul of 10% DMS in ethanol or ethanol alone at 37°C for 5 minutes followed by incubation with 20% beta-mercaptoethanol for 5 minutes on ice. Treatment with the chemical probe was followed immediately by RNA XP bead cleanup and error-prone reverse transcription (RT) using SuperScript™ II (Invitrogen, ThermoFisher Scientific) with modified buffer (10X: 500 mM Tris pH 8, 750 mM KCl). 6nM manganese, RNase inhibitor, and reverse primer 5’-TGTTGGAGTCACTCGACTCCGGT-3’ were added to the RT reaction. The RT product was cleaned up with RNA XP beads and used as the template for second strand synthesis (Invitrogen Second Strand cDNA Synthesis Kit; ThermoFisher Scientific). Libraries were prepared using the Nextera DNA Library Preparation Kit (Illumina). The paired-end libraries were sequenced at 2 x 150 on an Illumina NovaSeq. Experiments were performed in duplicate.

### Structure probing analysis

Structure probing analysis was conducted with ShapeMapper v2.2.0 (53) using default parameters and the “--dms” flag for DMS probing samples. The DMSO/EtOH reads were designated as “untreated” and the 5NIA/DMS reads as “modified”, respectively. The Pearson correlation between replicates was calculated (Supp Table S1), and due to highly similar reactivity profiles, the replicates were merged. Normalized reactivity profiles were limited between 0 and 4 and utilized for downstream analysis. Representative 2D structures for human and chicken sequences were generated with RNAstructure v6.5 (54) incorporating reactivity data and a max distance of 66 base pairs. Regions bound by primers for library prep were set as single-stranded.

### Pairing Probability Calculations

Partition files were made for all microexon and midexon sequences using RNAstructure v6.5 (54) partition command with a max distance of 66 base pairs (Figure_5/partition.sh). The ProbabilityPlot command was used to calculate base pairing probabilities (Figure_5/pairprob.sh) and a custom python script was used to sum the probabilities at each nucleotide position (Figure_5/pairprobsum.py).

### Sliding Reactivity similarity

Human and chicken sequence alignments from PairwiseAligner from the Bio.Align package with default parameters were used to align normalized reactivity values. Spearman correlations between human and chicken SHAPE reactivities were calculated for both the full sequences (Supp Table S6; Figure_2/spearman_reactivity_correlation.R) and centered 15-nt sliding windows (step size = 1). For each window, sequence identity was calculated as the number of matching nucleotides divided by the window size (15 nts). Positions containing missing values were included in the window, but windows with less than 8 valid reactivity values were skipped. To determine significance, both human and chicken reactivity values were shuffled, and the correlation calculations were repeated 1000 times for both the full sequences and centered 15-nt sliding windows (Figure_2/observed_struct_correlation_vs_shuffled.R). A null distribution of correlation run lengths were computed for each microexon with a run counting as 2 or more neighboring windows having Spearman correlations > 0.2. The number of runs at any given length were compiled across the 1000 iterations. The length at which < 5% of runs were longer was set as the threshold for significance. Runs longer than this threshold in the unshuffled data were interpreted to include regions of highly similar reactivity profiles (Figure_2/identify_struct_similar_regions_agap1.R; Figure_2/sequence_identity_struct_overlay_agap1.R). All correlations and plots were done in R v4.1.2.

## Results

### Neural microexons are dynamically regulated during chick embryonic brain development

Human embryonic tissue and organoid models suggest that microexons are dynamically regulated during brain development, however, these samples are limited in accessibility and may not be representative of normal human development (55,56). We employed the chick embryo model system to analyze microexon splicing during embryonic development. We dissected embryonic brain and heart tissue at five time-points (Hamburger-Hamilton Stages 11, 15, 27, 31 and 35), prepared sequencing libraries and performed short read sequencing. We identified 347 microexons with distinct inclusion and exclusion patterns in brain tissue (Supp Figure 1). We focused on 13 selected neural microexons that are highly expressed, conserved between human and chicken and are associated with human neurological disorders (Table 1). Looking at this subset of neural microexons, the transition between early stages HH11-15 and later stages HH27-35 is especially dramatic, with nearly half of selected microexons sharply increasing in inclusion at this developmental stage in the brain (Figure 1A; *FRY microtubule binding protein (FRY), AGAP1, protein tyrosine phosphatase receptor type K (PTPRK), CPEB4, amyloid beta precursor protein binding family B member 2 (APBB2)* and *DCTN1*). The inclusion of the *ROBO1* microexon decreases at the transition between HH15 and HH27, consistent with its known biological role in axon guidance (9) (Figure 1A). Only two microexons remain relatively stable at this transition point in chicken brain tissues (Figure 1A, dedicator of cytokinesis 7 (*DOCK7)* and myocyte enhancer factor 2A (*MEF2A))*. Neither sex nor transcript expression levels are correlated with microexon splicing (Supp Figure S2-4). This regulated splicing between HH15 and HH27 is representative of broader expression changes in chicken neural microexons (Supp Figure S1).

**Figure 1.**
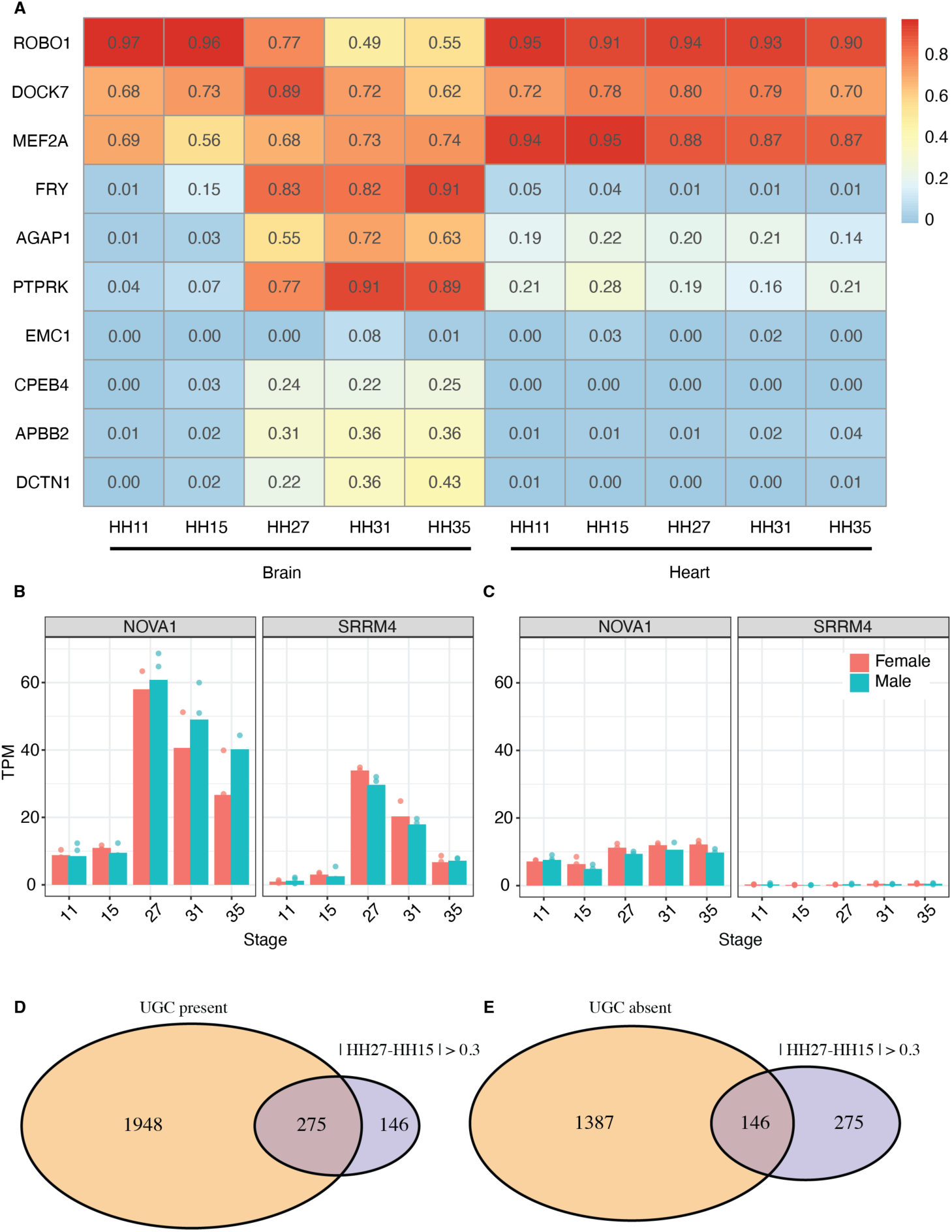
The majority of select neural microexons are spliced in a tissue-specific and temporally regulated manner during chick embryonic development. (A) Heatmap of select chicken neural microexons PSI values during HH11, HH15, HH27, HH31, and HH35 developmental stages of brain and heart tissue. Includes triplicate male and female chickens. (B) Expression of the RNA binding proteins *NOVA1* and *SRRM4* during brain and (C) heart development. (D) Overlap of microexons containing at least one UGC motif in the branchpoint-to-splice site region with microexons containing PSI +/- 0.3 between HH15 and HH27 stages. (E) Overlap of microexons containing no UGC motifs in the branchpoint-to-splice site region with microexons with PSI +/- 0.3 between HH15 and HH27 stages.

**Table 1.** Selected conserved neural microexons and their properties (37). Gene function summarized from (3,56,58)

| *Gene* | *Human Expression (RPKM)* | *Human Average PSI (%)* | *Gene Function* | *Microexon Function* | *Human to Chicken Sequence Alignment* | *Covariation (9 model / 100 vertebrates)* |
| --- | --- | --- | --- | --- | --- | --- |
| *AGAP1* | 15.8 | 13.1% | Endolysosomal trafficking | unknown | 79.3% | yes/yes |
| *APBB2* | 10.4 | 14.8% | Amyloidogenesis | unknown | 67.5% | yes/yes |
| *ASAP2* | 14.7 | 8.3% | Cellular migration | unknown | 72.0% | yes/no |
| *CLEC16A* | 6.4 | 9.7% | Autophagy and mitophagy regulation | unknown | 72.0% | no/no |
| *CPEB4* | 15.9 | 12.4% | Polyadenylation regulator | CPEB4 condensate interactions (6,7) | 67.8% | yes/no |
| *DCTN1* | 76.1 | 11.9% | Axonal transport | unknown | 62.5% | no/no |
| *DOCK7* | 8.3 | 19.8% | Neuronal polarization | unknown | 76.5% | yes/yes |
| *EMC1* | 11.4 | 13.9% | Membrane protein biogenesis | unknown | 66.5% | yes/yes |
| *FRY* | 10.3 | 21.3% | Bipolar spindle organization | unknown | 71.3% | yes/no |
| *ITSN1* | 9.8 | 6.4% | Signaling | Tissue specific protein interactions (59) | 68.8% | yes/no |
| *MEF2A* | 17.8 | 13.6% | Neuronal differentiation and survival | unknown | 69.5% | yes/yes |
| *PTPRK* | 11.7 | 11.2% | Synaptic transmission | unknown | 69.5% | yes/yes |
| *ROBO1* | 27.7 | 88.5% | Axon guidance | Midline crossing (9) | 79.3% | yes/no |

At the transition point between HH15 and HH27 where we identified dynamic microexon regulation (Figure 1A), we also see dramatic increases in expression of the RNA binding proteins *NOVA1* and *SRRM4* in the brain at the RNA level, followed by a slow decrease from HH27 to HH31 and HH35 (Figure 1B). NOVA1 and SRRM4 are well-known microexon regulatory factors that influence human microexon splicing (5). *SRRM4* is expressed 50-fold higher in brain and *NOVA1* is expressed 6-fold higher in brain than in heart tissue (Figure 1B and C; Supp Figure S5). Two other RNA binding proteins may also be relevant for neural specific splicing, *RNA binding fox-1 homolog 1 (RBFOX1)* and *RNA binding fox-1 homolog 3 (RBFOX3),* as they also have higher expression in brain than in heart tissue and increase at the HH27 transition (Supp Figure S5). Other splicing factors linked to microexon regulation, *polypyrimidine tract binding protein 1 (PTBP1), serine and arginine rich splicing factor 11 (SRSF11)* and *RNA binding protein with serine rich domain 1 (RNPS1)*, have similar RNA expression in both brain tissues and heart tissue and may not be specific for neural microexon regulation (Supp Figure S5). SRRM4 is associated with UGC motifs in microexons to regulate their splicing (23). Microexons with large changes in splicing at the HH15-HH27 transition are more likely to contain UGC motifs in the 3’ splice site region (Figure 1D). However, a significant portion of dynamic brain-associated microexons do not have UGC motifs consistent with SRRM4 (Figure 1E).

### Neural microexon conserved RNA structures correspond to regions of high sequence similarity

To analyze chicken and human microexons regulatory motifs across species, we developed a list of conserved human and chicken neural microexons. We classified microexons into neural and non-neural groups using the VastDB splicing database (37). Consistent with prior studies, we found that neural microexons and their proximate introns had high conservation compared to non-neural microexons or moderately sized exons (Supp Figure S6-9) (3,4). We also confirmed that introns flanking neural microexons are significantly shorter than introns flanking midexons in humans (2,27) (Supp Figure S10A-B), but this was not true for chickens introns (Supp Figure 10C-D). Interestingly, we found that introns flanking human neural microexons have significantly less repetitive sequence than introns flanking midexons (Supp Figure S10E-F, Supp Figure 11), supporting a functional role for introns flanking microexons.

RNA structure is known to influence precursor RNA processing, but little is known about the RNA structure around microexons in precursor mRNAs (12). We analyzed the structure of our 13 selected microexons (Table 1). We expected RNA structures to be highly similar between humans and chickens, as RNA structure can maintain more information about functional conservation than sequence alone (50,57). Covariation analysis can detect nucleotide variation that preserves base pairing and thus maintains functional RNA structures (50). We performed covariation and conservation analysis using alignments at our 13 select microexons from 9 model species and 100 vertebrates. We found evidence for covariation at 11 microexons using model species and at 6 microexons using 100 vertebrates (Table 1). For example, conservation and covariation analysis of a microexon in *ArfGAP with GTPase domain, ankyrin repeat and PH domain 1* (*AGAP1*) suggests the presence of a stem-loop encompassing the microexon (Figure 2A).

**Figure 2.**
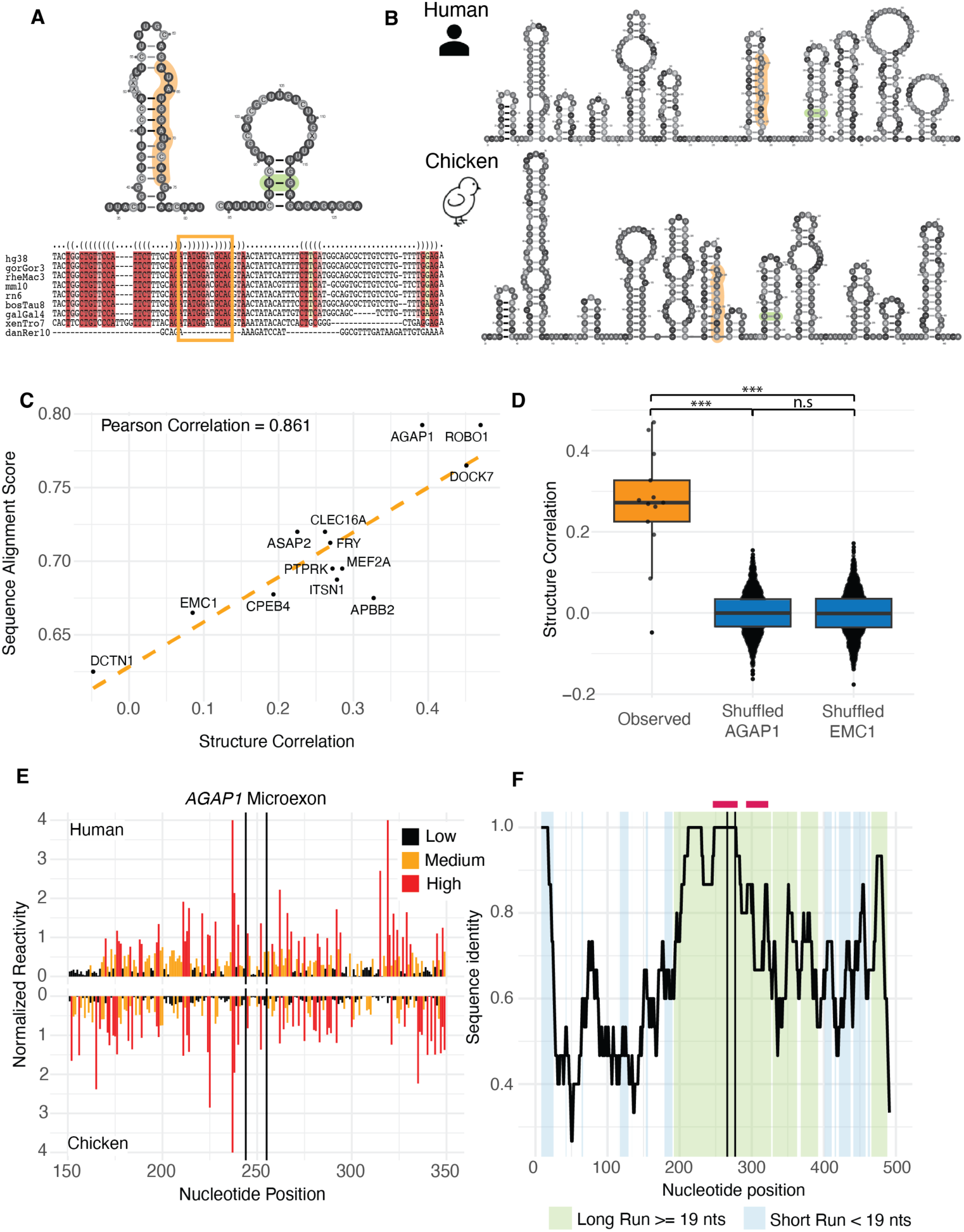
Secondary structure depends on sequence conservation in *AGAP1* and other microexons. (A) Predicted *AGAP1* structure and sequence alignment based on covariation (green) and conservation (red) within common model organisms. Microexon (orange) and covariate positions (green) highlighted. (B) Representative structures of human (top) and chicken (bottom) *AGAP1*. (C) Correlation of sequence similarity and structural similarity of select microexons (Spearman). (D) Structural similarity between human and chicken compared to background (Spearman correlation, background =1000 iterations of shuffled reactivity values, Wilcoxon signed-rank test, two-sided; *= p-value < 0.05, ** = p-value < 0.01, *** = p-value < 0.005, n.s = p-value > 0.05). (E) Reactivity profiles of human (top) and chicken (bottom) *AGAP1*. (F) *AGAP1* sequence identity across aligned human and chicken sequences using a 15-nt sliding window (step size = 1nt). Green or blue shading indicates regions >= 19 nts or < 19 nts, respectively, with Spearman correlations between aligned human and chicken reactivities of >= 0.2; Red bars indicate position of structures predicted with covariation and conservation data. Splice sites indicated by black vertical lines.

We then developed experimentally-based *in vitro* structural models of all 13 microexons and local intronic regions in both human and chicken precursor mRNAs. We performed selective 2’ hydroxyl acylation by primer extension (SHAPE) and mutational profiling (SHAPE-MaP) and dimethyl sulfide (DMS) mutational profiling (DMS-MaP) chemical probing and found high reproducibility between replicates (∼0.9). We also found reasonable correlation between SHAPE and DMS data, (∼0.4) (Supp Tables S1 and S2) (51,52). To confirm that *in vitro* structure data was representative of the RNA structure in the cell, nuclei from a medulloblastoma cell line were isolated and probed with both 5NIA and DMS. *In vitro* and cellular SHAPE reactivities were highly correlated (mean ∼0.7) (Supp Figures S12-15, Supp Table S5). *In vitro* and cellular DMS-MaP experiments were less correlated (∼0.5), but less accurate as DMS provides highly accurate reactivity primarily for adenines and cytosines, which were limited in our polypyrimidine tract-rich cellular targets (Supp Table S3).

In *AGAP1*, we found that the nucleotides involved in the predicted covariation are maintained as base-pairs in a stem-loop downstream of the microexon in representative structures for both human and chicken (Figure 2B). However, predicted covarying nucleotides were not consistently found in the secondary structure predictions in the other 10 microexons containing potential covariation (Supp Figures S16-27). In addition, base-pairing predicted by covariation/conservation did not successfully predict low SHAPE or DMS reactivity overall (Supp Figure S28). We do not find evidence for covariation as a mechanism to maintain conserved RNA structures around neural microexons. Instead, we find that orthologous sequences have structural similarity dependent on sequence similarity (Figure 2C, Supp Table S6). For example, the high sequence similarity between humans and chickens in *AGAP1* (0.79) corresponds to high structural correlation between humans and chicken based on experimental reactivity data (0.39). This structural similarity is significantly higher than randomized reactivities (Figure 2D), but not suggestive of independent conservation of structure. Interestingly, despite relatively high sequence similarity scores (>0.5), the microexons in ER membrane protein complex subunit 1 (*EMC1*) and dynactin subunit 1 (*DCTN1*) could not be distinguished from background as structurally similar across species (Figure 2C). Our data do not suggest that microexons or their surrounding intronic sequence maintain high levels of RNA structural similarity independent of sequence conservation.

We observed that human and chicken sequences frequently contain small insertions or deletions, which could contribute to low structure similarity scores. Therefore, we focused on regions of contiguous high similarity scores to identify whether small local regions of structure are conserved between humans and chickens. We identified blocks of structural similarity and found that these blocks typically include seeds of sequence similarity of at least 0.5 (Figure 2F; Supp Figures S16-27). *AGAP1* contains extensive structural blocks encompassing the microexon and region of predicted covariation/conservation (Figure 2F). This suggests that small regions of structure in flanking introns and microexons are conserved between humans and chickens. However, these structural blocks are not located in a consistent pattern within the 13 tested microexons and can be upstream, downstream or in the microexon (Figure 2F, Supp Figures S16-27). Our data suggest that neural microexons do not have high levels of structural conservation or broad global structural conservation but have maintained blocks of structural similarity that primarily correlate with high sequence similarity. Due to their varied localization and length within our selected microexons, these regions of structural similarity likely contribute in unique ways to the regulation of each individual precursor RNA.

### The branchpoint-to-splice site region in neural microexons is lengthened by an extended polypyrimidine tract and structurally accessible nucleotides

Due to their small size, microexons have been found to require robust splicing sequence signals to promote exon inclusion (2). Splicing sequence signals can be compared to 5’ and 3’ splice site consensus sequences and quantified for strength (MAXENT score, (41)). As expected based on prior work (3), we found that the 3’ splice sites of neural microexons were significantly weaker than the 3’ splice sites of neural midexons in humans (Figure 3A and B) and in chickens (Supp Figure S29A). Both human midexons and non-neural microexons have higher polypyrimidine frequencies between 5 to 15 nts from the 3’ splice site, but we found that neural microexons have the highest polypyrimidine frequency between 15 to 30 nts from the GU (Figure 3C, Supp Figure S29C). This extension of the polypyrimidine tract is not apparent at the downstream constitutive 3’ splice site for neural microexons, midexons, or non-neural microexons in humans or chickens (Figure 3D, Supp Figure S29D).

**Figure 3.**
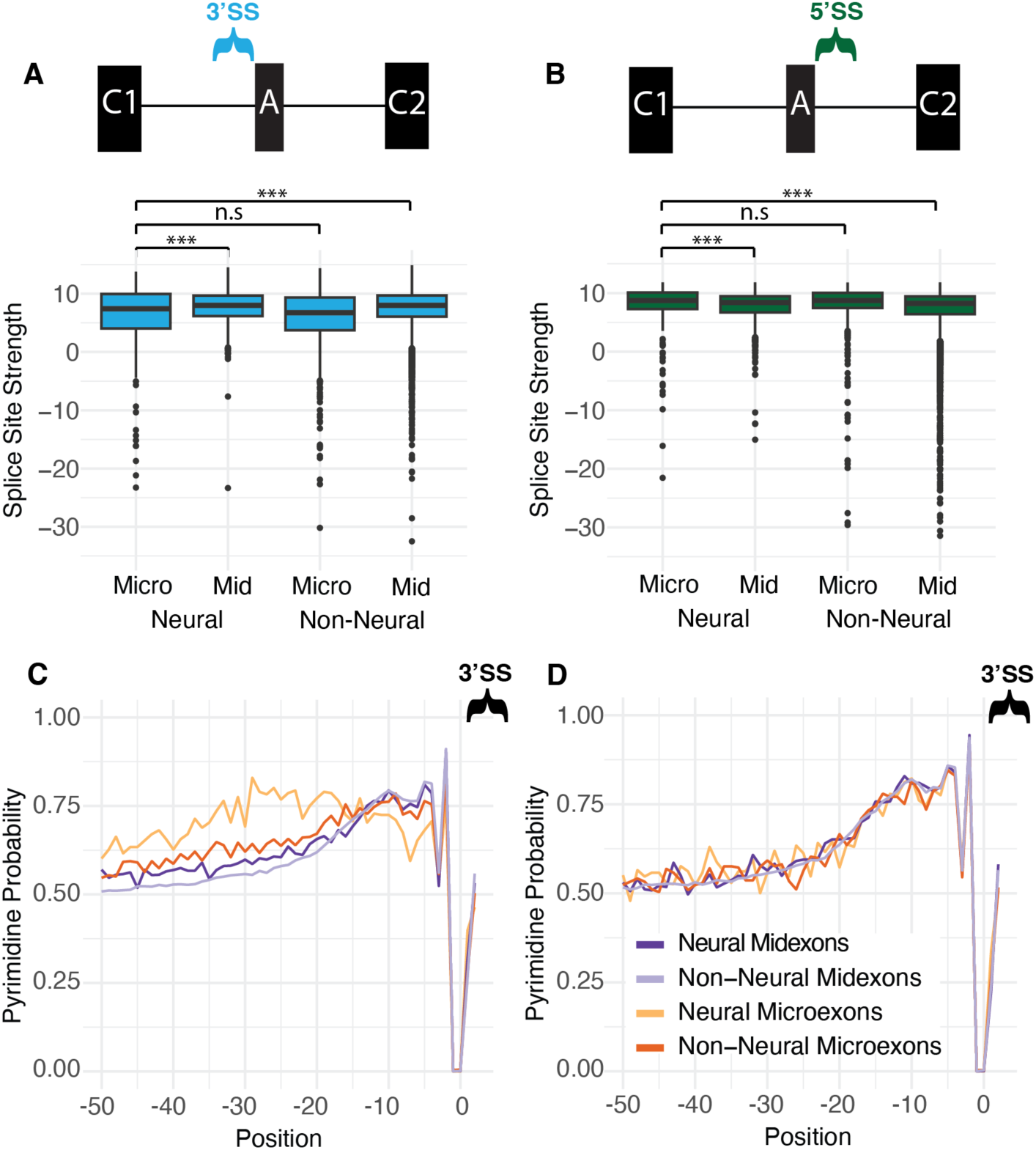
Human neural microexons have weak 3’ splice sites and a shifted polypyrimidine tract. (A) 3’ splice site strength scores for human microexon and midexon sequences. (B) 5’ splice site strength scores for human microexon and midexon sequences. (C) Probability of cytosine or uracil per nucleotide position upstream of alternatively microexons and midexons. (D) Probability of cytosine or uridine per nucleotide position approaching the 3’ss of the exons downstream of the alternatively spliced microexons and midexons. (human: neural midexons n=1053, non-neural midexons n=21787, neural microexons n=276, non-neural microexons n=554; chicken: neural midexons n=1700, non-neural midexons n=8210, neural microexons n=249, non-neural microexons n=403; Wilcoxon signed-rank test, two-sided; * = p-value < 0.05, ** = p-value < 0.01, *** = p-value < 0.005).

Based on the unusual properties of polypyrimidine tracts in neural microexons (Figure 3C), we analyzed the predicted location of the branchpoint using SVM-BPfinder (Corvelo et al. 2010). We found, in both humans and in chickens, that the branchpoint was predicted to be significantly farther from the 3’ splice site around neural microexons, (median of 45 vs 34 nts in humans, 47 vs 34 nts in chicken, Figure 4A and B). This is consistent with observations of extended 3’ splice site regions containing U/C and UGC SRRM4 binding sites in autism-associated microexons (23). Unsurprisingly, insertion of UGC motifs results in longer branchpoint-to-splice site distances (Figure 4C and D) and polypyrimidine tract lengths in both microexons and midexons in humans and chickens (Supp Figure S30). In chickens the branchpoint-to-splice site length of neural microexons are significantly longer than that of neural midexons (Figure 4D), and this trend is non-significant but consistent in humans (Figure 4C).

**Figure 4.**
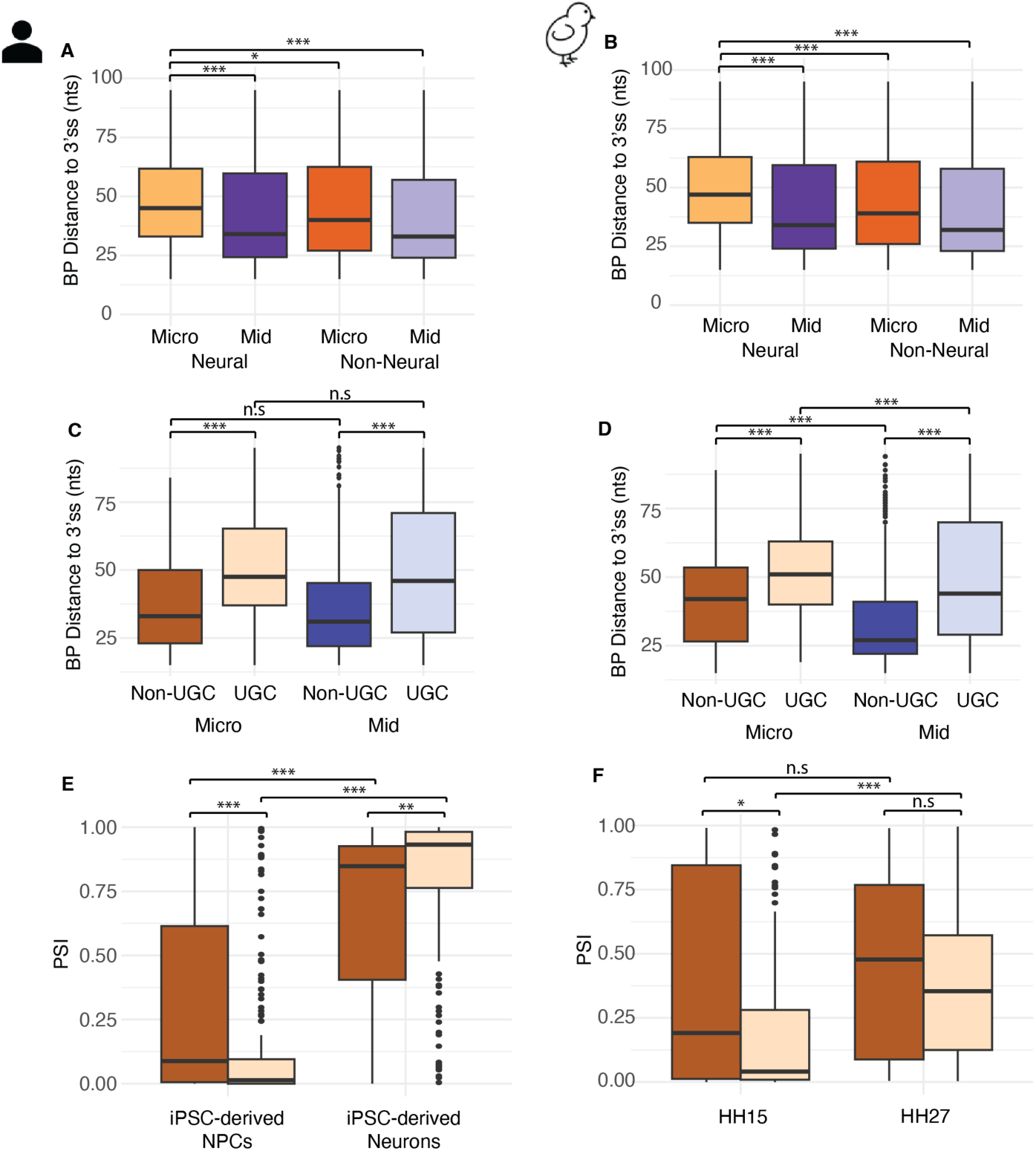
Neural microexons have an extended branchpoint-to-splice site region through unique sequence elements associated with increased splicing efficiency during neurodevelopment. (A) Distance between the top-scoring branchpoint (BP) and the 3’ splice site of human and (B) chicken alternative exons. (C) Distance between the top-scoring branchpoint and the 3’ splice site of neural UGC containing and non-UGC containing microexon and midexons in human and (D) chicken. (E) PSI of neural UGC and non-UGC containing microexons in human iPSC-derived NPCs and iPSC-derived neurons. (F) PSI of neural UGC and non-UGC containing microexons in chicken HH15 and HH27 brains. (human: non-UCG microexons n=51, UGC microexons n=180, non-UGC midexons n=338, UGC midexons n=353; chicken: non-UCG microexons n=39, UGC microexons n=117, non-UGC midexons n=469, UGC midexons n=585; neural microexons n = 13, neural midexons n = 7, Wilcoxon signed-rank test, two-sided; *= p-value < 0.05, ** = p-value < 0.01, *** = p-value < 0.005, n.s = p-value > 0.05).

Both UGC and non-UGC containing neural microexons increase dramatically in inclusion during development and differentiation. After differentiation from human induced pluripotent stem cell-derived (hiPSC) neural precursors to mature neurons, both UGC and non-UGC containing microexons significantly increase in inclusion (36). Likewise, at the transition point from HH15 to HH27 in chick development, we see a significant increase in UGC and a non-significant increase in non-UGC containing microexons (Figure 4E and F). This suggests one mechanism of circumventing the constraints of such small exon length is extension of the branchpoint-to-3’ splice site region to provide physical space, regardless of UGC motifs, which may recruit SRRM4 and other splicing factors to promote exon inclusion.

In addition, using our experimental reactivity data from our selected microexons, we find that the human branchpoint-to-3’ splice site region has significantly higher structural accessibility compared to experimental reactivity data from a set of human neural midexons (Figure 5A). The high reactivity in the branchpoint-to-splice site region indicates that this region is accessible and likely extended. In contrast, we did not find any other patterns of consistent accessibility in our experimental reactivity data at core elements, including the branchpoint, 3’ splice site and 5’ splice site (Supp Figure S31). These analyses are based on experimental data from a small set of microexons, however, computational prediction shows that both human and chicken microexons are highly accessible in the branchpoint-to-3’splice site region (Figure 5B and C). This suggests that neural microexons use a distinct and shared splicing mechanism that lengthens the distance between the branchpoint and splice site through both sequence and structural elements, potentially allowing for exon-definition to assist in splice site identification.

**Figure 5.**
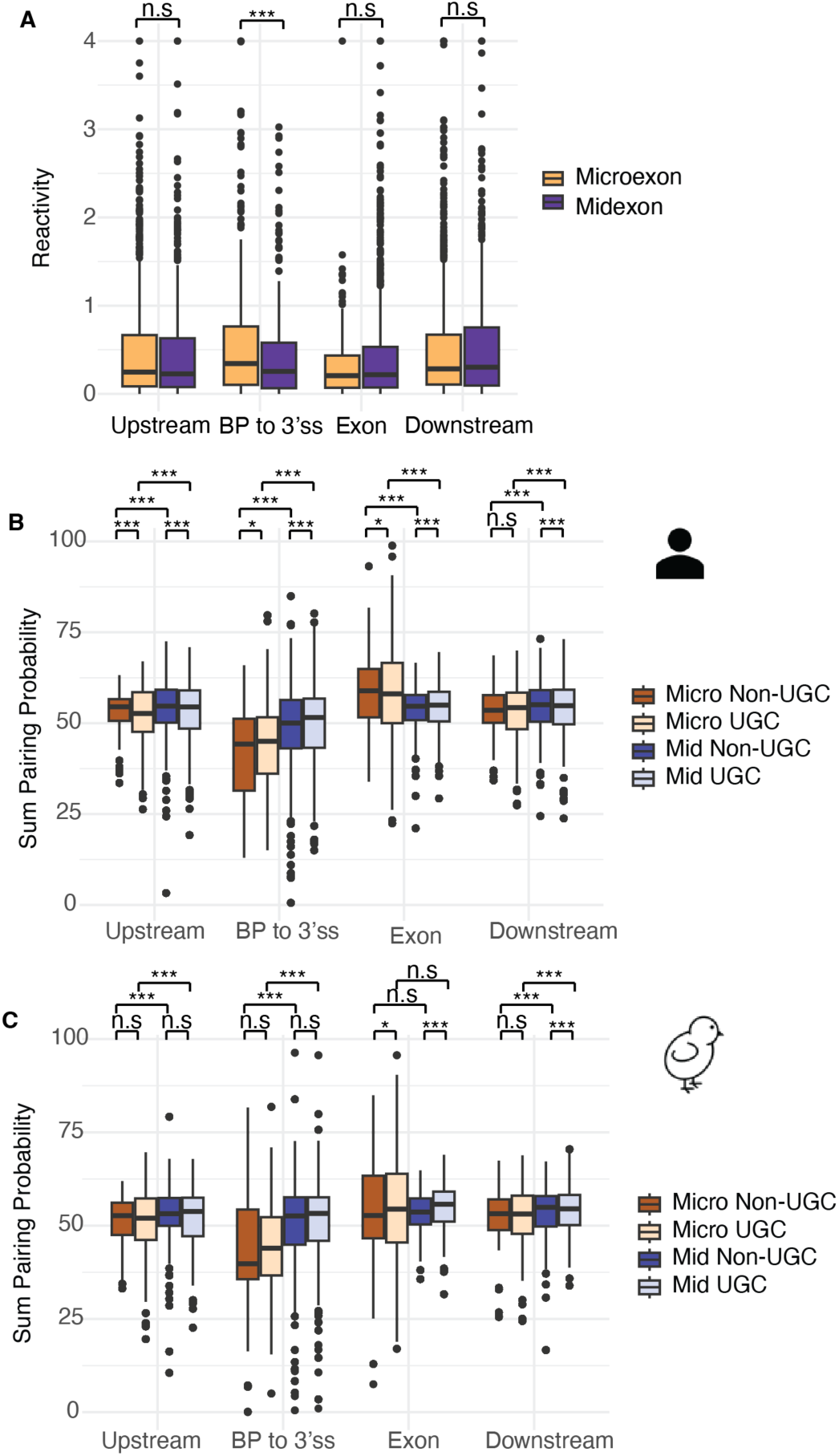
Neural microexons have an extended branchpoint-to-splice site region through unique RNA structure elements. (A) Experimental nucleotide reactivities at different regions of human microexons and midexons, including 100nts upstream of the branchpoint (BP), between the BP and 3’ splice site, within the exon, and 100nts downstream of the 5’ splice site (neural microexons n=13, neural midexons n=7). (B) Computational mean summed pairing probability of nucleotides in human and (C) chicken neural UGC containing and non-UGC containing microexons and neural midexons at the same regions. (human: non-UCG microexons n=51, UGC microexons n=180, non-UGC midexons n=338, UGC midexons n=353; chicken: non-UCG microexons n=39, UGC microexons n=117, non-UGC midexons n=469, UGC midexons n=585; Wilcoxon signed-rank test, two-sided; *= p-value < 0.05, ** = p-value < 0.01, *** = p-value < 0.005, n.s = p-value > 0.05).

## Discussion

Microexons present a conceptual challenge for canonical models of splice site recognition due to their extremely small size. The exon-definition model, which relies on coordinated recognition of splice sites across an exon, is incompatible with microexons because they lack sufficient sequence length to physically accommodate cross-exon interactions (16). In this study, we identify factors that contribute to lengthening of the crucial branchpoint-to-splice site region including 1) distal placement of the polypyrimidine tract farther into the intron and 2) extension of the polypyrimidine tract sequence. In addition, we find that 3) the branchpoint-to-splice site region is highly accessible and likely unpaired in neural microexons.

Due to shared sequence content, we were surprised to find little evidence that neural microexons rely on conserved RNA secondary structures maintained through covariation. Although covariation analysis predicted conserved base-pairing around many of the selected microexons, these predictions were not supported by experimental SHAPE reactivity data. Moreover, structural similarity between orthologous human and chicken precursor RNAs was relatively low, even in cases of high sequence similarity. The lack of strong structural or accessibility conservation suggests that, for neural microexons, global structure is not strictly conserved for function across species. We find evidence for local conserved structures at unique positions in each neural microexon we analyzed. These local structures, in blocks of conserved sequence, may be adequate for functional regulation. In addition, other structural properties, such as decreased pairing within the extended branchpoint-to-splice site region, may be critical instead of specific structural motifs. Future studies will be important for directly testing the mechanistic role of the extended branchpoint-to-splice site region in splicing efficiency.

We are interested in functional roles for neural microexons; however, many of these genes are difficult to study as they are involved in embryonic brain development (3,56). The chick embryo is a well-established developmental model system (28). We harvested brain and heart tissue from five embryonic developmental stages. We find that the majority of our selected neural microexons undergo dynamic, stage-specific regulation during brain development. The timing of increased microexon inclusion coincides with elevated expression of known microexon regulators, including SRRM4 and NOVA1, supporting a conserved regulatory program across vertebrates. Microexon splicing is regulated at the splicing level during brain development and not with transcriptional changes to the microexon gene as changes in microexon inclusion/exclusion are not correlated with gene expression. The chick embryo is an accessible developmental system that allows for perturbation of regulation, including splicing, and analysis of downstream impact on morphology and function. Future studies on the role of broad changes to microexon inclusion, such as through SRRM4 knock-down, and targeted changes to specific microexons, such as microexon exclusion within *ROBO1*, are exciting potential future experiments with this model system.

Limitations of this study should be acknowledged. Branchpoint positions were inferred computationally using a program based on mammalian splicing (45). Although human and chicken ribosomal RNA sequences are highly similar, and consistent trends were observed across both species, branchpoint prediction may be less accurate for chickens. In addition, our structural assays were performed *in vitro*. However, we find high correlation between our *in vitro* results and the microexon RNA structure derived from cellular RNA, suggesting that the *in vitro* RNA structure is representative of *in vivo* RNA. However, these experiments may not fully capture the influence of co-transcriptional folding, RNA-binding proteins, or spliceosomal components.

In summary, we find that neural microexons do not share specific local structures between orthologous regions in human and chicken but have unique regions of similar structural accessibility that correlate with high sequence similarity. Both human and chicken neural microexons have more accessible branchpoint-to-splice site regions than moderately-sized exons. These features suggest a mechanism by which neural microexons may overcome size constraints, such as steric hinderance for spliceosomal proteins or regulatory proteins like SRRM4. This work provides a foundation for future mechanistic studies of microexon splicing regulation.

## Data availability statement

Scripts used for data analysis and plotting are available at https://github.com/randazzal/extended_bp_2_3ss_scripts. Raw sequencing reads from embryonic chicken heart and brain tissues will be deposited to the gene expression omnibus. Other data available upon request.

## Acknowledgments

The authors want to acknowledge Dr. Susan Chapman for her valuable mentorship, training, and equipment sharing which facilitated the chicken embryo work in this manuscript. In addition, the authors wish to acknowledge the Institute for Human Genetics Molecular Core and the Institute for Human Genetics Bioinformatics Core, which supported the generation of the sequencing libraries and analysis presented in this work.

## Author Contributions

Conceptualization, A.R., L.L.; resources, A.R., K.E.H., J.R.M., A.H., T.D.O.-H., L.L.; data curation, A.R.; methodology, A.R., T.D.O.-H., L.L.; investigation, A.R., K.E.H., J.R.M., A.H.; formal analysis, A.R.; supervision, T.D.O.-H., L.L.; funding acquisition, T.D.O.-H., L.L.; validation, A.R., K.E.H., J.R.M., A.H.; visualization, A.R.; writing – original draft, A.R., L.L.; writing – review & editing, A.R., K.E.H., J.R.M., T.D.O.-H., L.L.; project administration, L.L.

