## Supplementary Material for "Conservation of extended sequence and structure in the branchpoint-to-3’ splice site region upstream of neural microexons"

- Supplemental Figures S1-31 ........................................ 2-32
- Supplemental Tables S1-5 ........................................ 33-34
- Supplemental Methods ........................................ 35-36

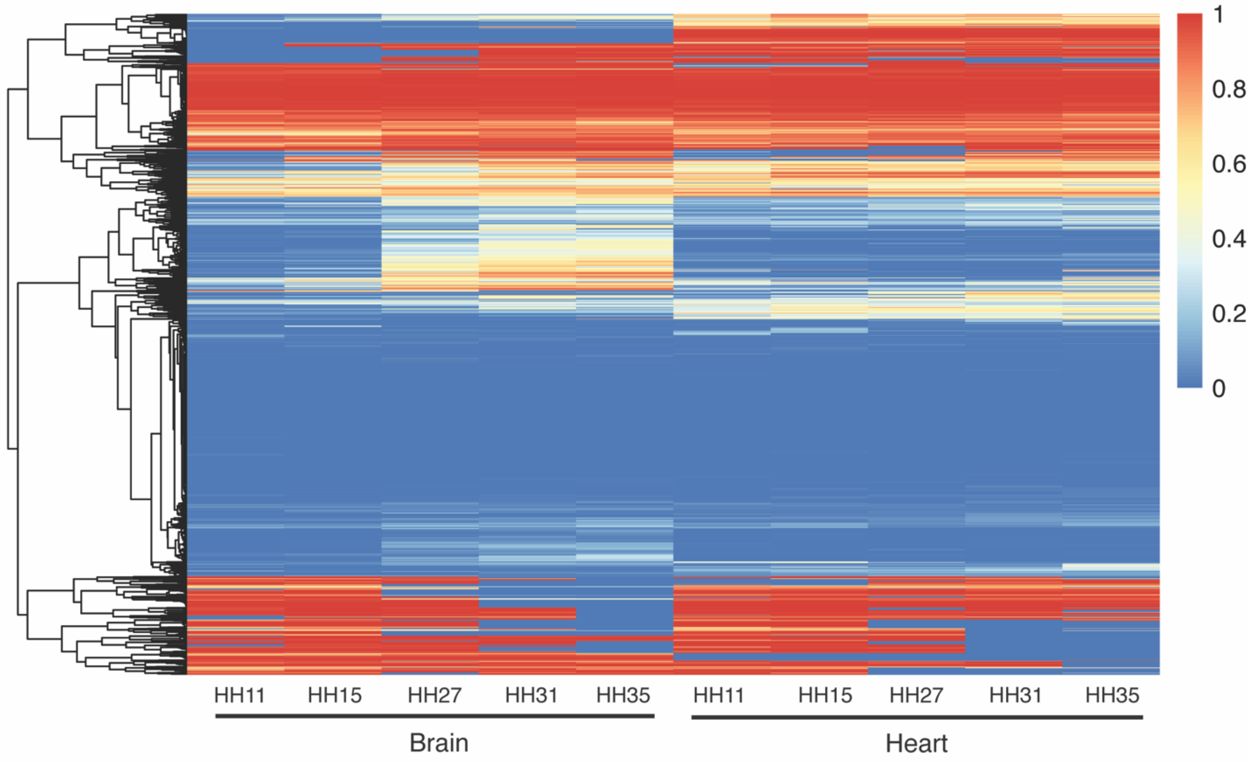


**Figure S1. Microexons exhibit different splicing patterns in neural and cardiac tissue development.** Heatmap of PSI values of microexons showing distinct splicing patterns during embryonic chicken brain and heart development.

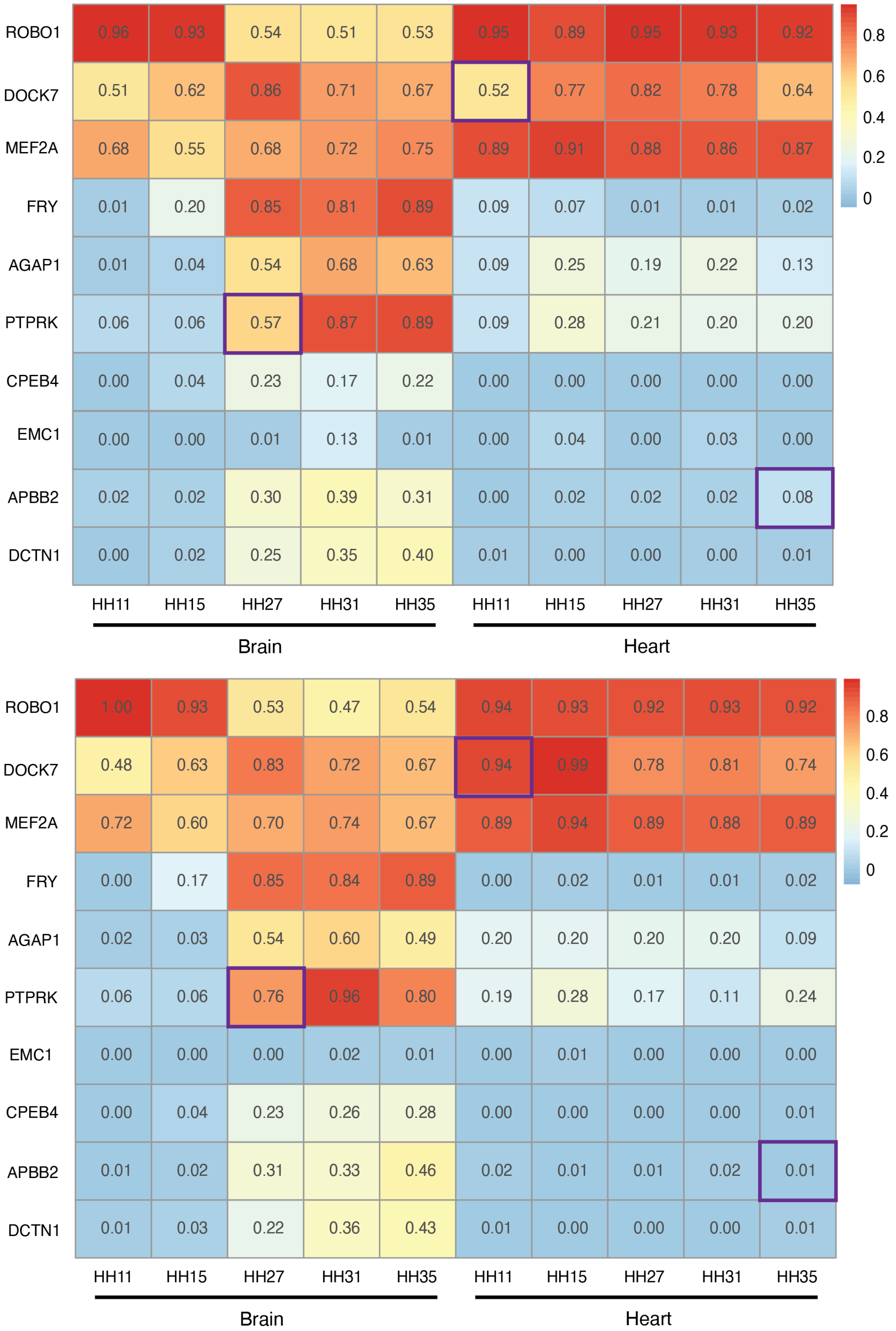


**Figure S2. Microexons are similarly spliced in male (top) and female (bottom) chickens.** Values indicate the average PSI level among replicates. Purple boxes indicate stages with significantly different PSI values between male and female chickens (FDR < 0.05).

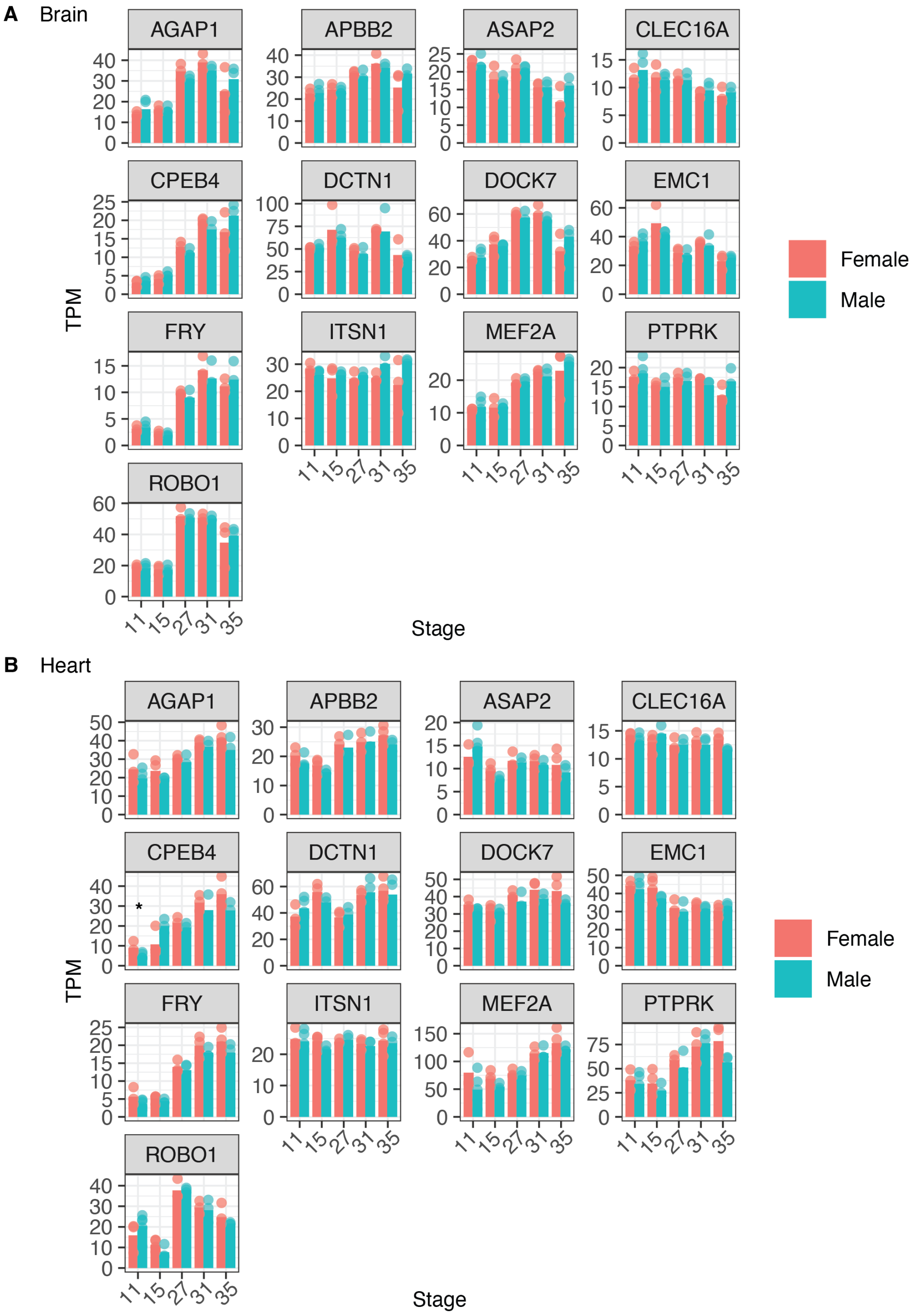


**Figure S3. Expression of microexon-containing genes undergoes tissue and temporal regulation shows little difference between sexes.** TPM values of select microexon-containing genes during chicken (A) brain and (B) heart development. (* = FDR < 0.05 between males and females).

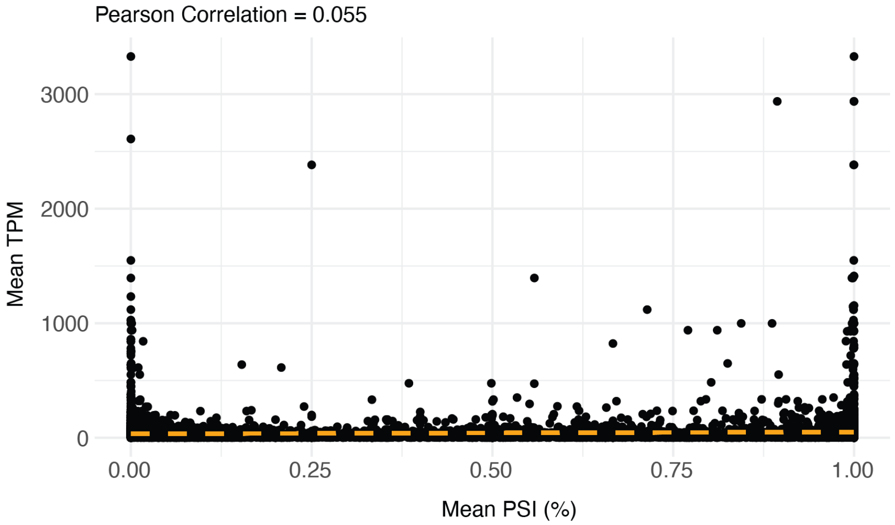


**Figure S4. There is no correlation between a microexon gene expression and microexon splicing inclusion.** Dot plot of mean TPM values and mean PSI values across replicates of both sexes. The dashed orange line indicates a linear regression fit, and the correlation was calculated with Pearson.

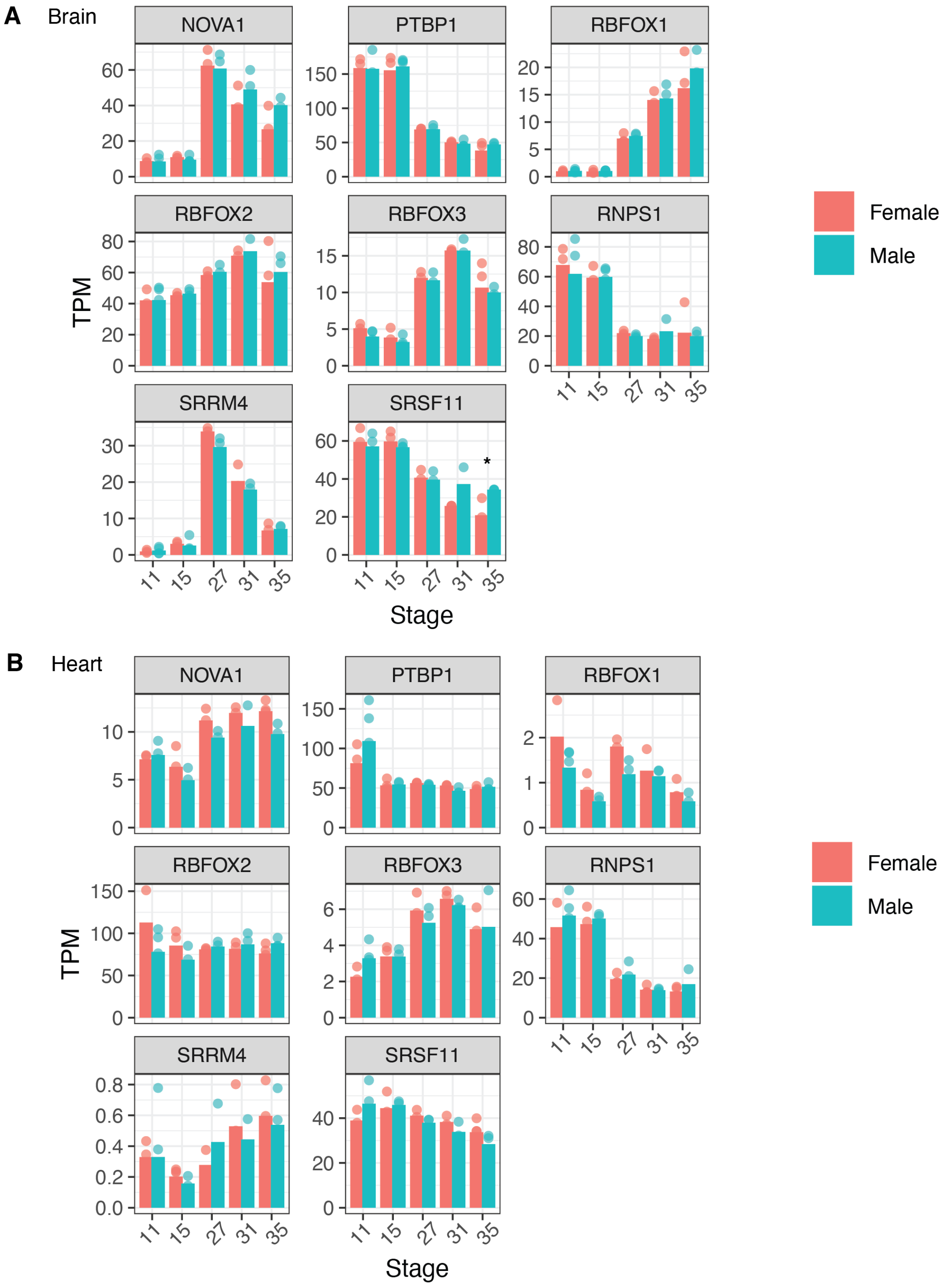


**Figure S5. Expression of microexon-associated RBPs undergoes tissue and temporal regulation.** TPM values of RBPs associated with microexon splicing regulation during chicken (A) brain and (B) heart development. Tested RBPs show little difference between sexes (* = FDR < 0.05 between males and females).

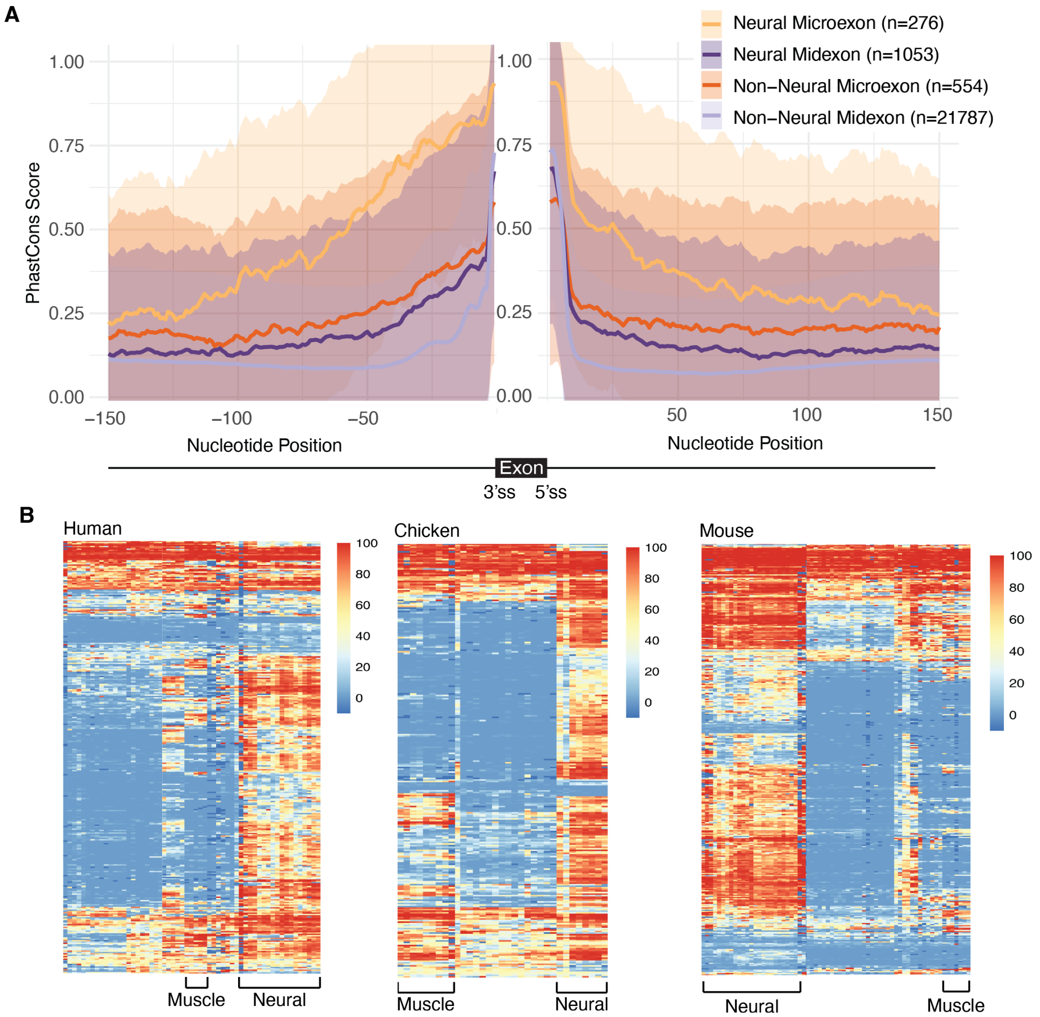


**Figure S6. Neural microexons are flanked by highly conserved intronic sequences and spliced in a tissue-specific manner.** (A) Mean phastCons scores across +/- 150 nucleotides of intronic sequence. Shaded regions show +/- 1 standard deviation. (B) Heatmap of microexon percent spliced in (PSI) values for a subset of human, chicken, and mouse tissues from VastDB. Dark blue indicates missing values. 3'ss = 3' splice site, 5'ss = 5' splice site.

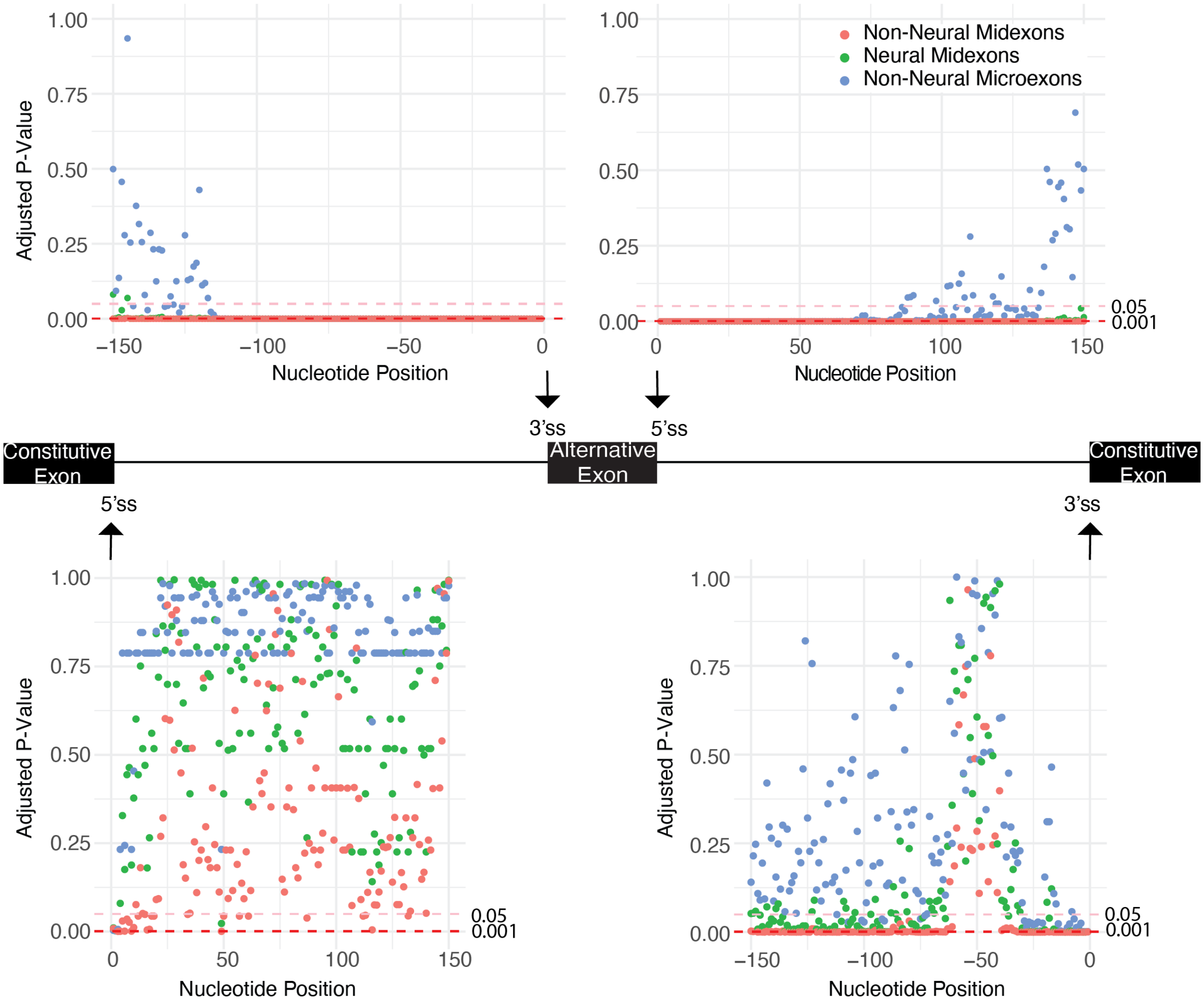


**Figure S7. Statistical significance for comparison of phastCons scores with neural microexons.** FDR adjusted p-values (Wilcoxon rank-sum test) at every nucleotide position approaching splice sites flanking microexons in Supplemental Figure S6A (top) and flanking constitutive exons in Supplemental Figure S8A (bottom).

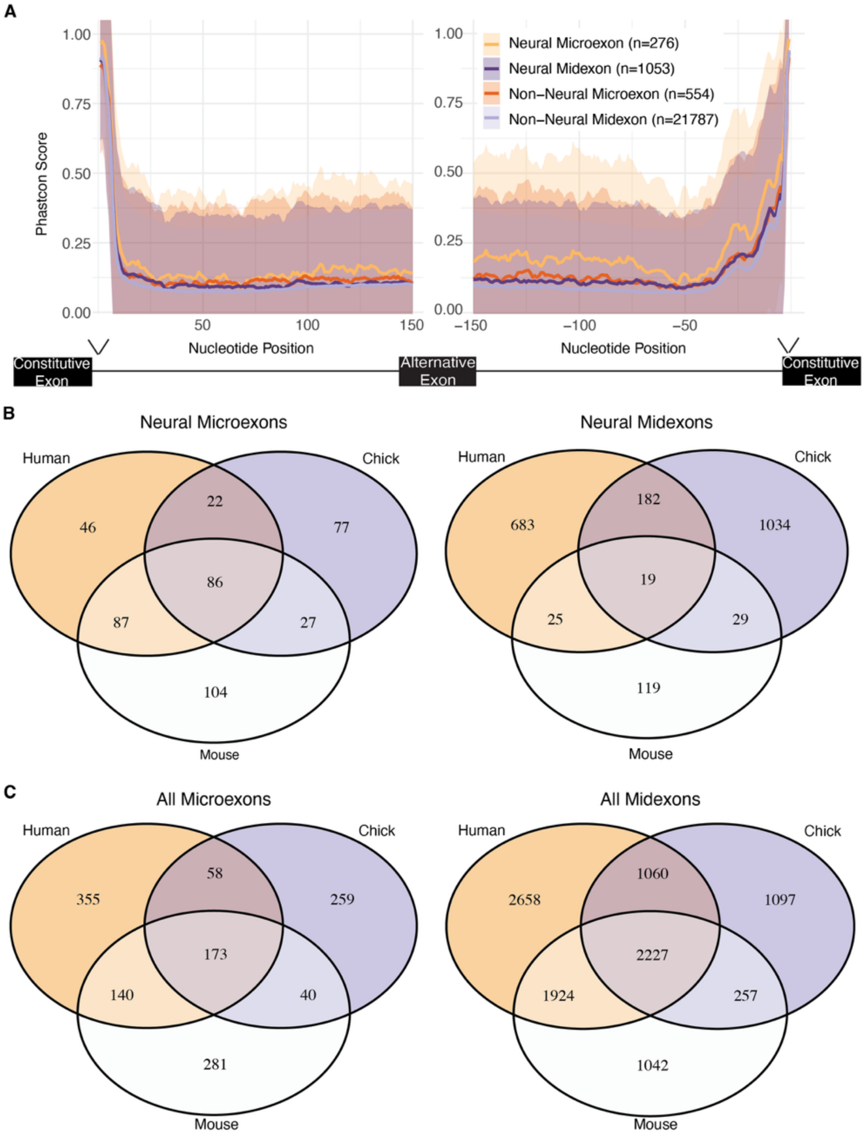


**Figure S8. Neural Microexons are highly conserved between species.** (A) Mean phastCons score approaching constitutive exons flanking the selected alternative microexons and midexons with shading showing +/- 1 standard deviation from the mean. FDR adjusted p-values per nucleotide position in Supplemental Figure S7. (B) Overlap of genes in human, chicken, and mouse containing neural microexons (left) or neural midexons (right). (C) Overlap of genes in human, chicken, and mouse containing microexons (left) or midexons (right).

A

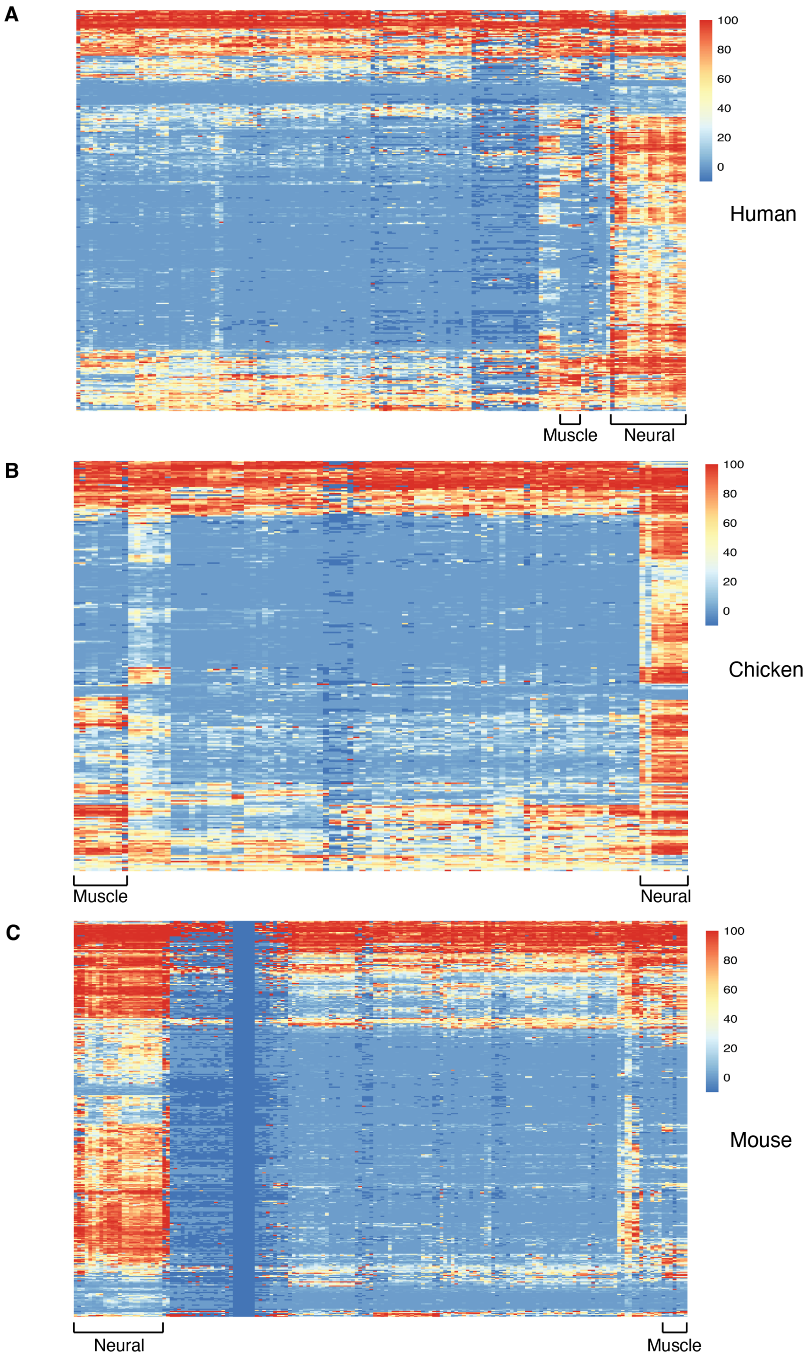


**Figure S9. A subset of microexons is spliced in a tissue specific manner.** (A) Heatmap of human microexon PSI values in all tissues. (B) Heatmap of chicken microexon PSI values in all tissues. (C) Heatmap of mouse microexon PSI values in all tissues. Dark blue indicates missing values.

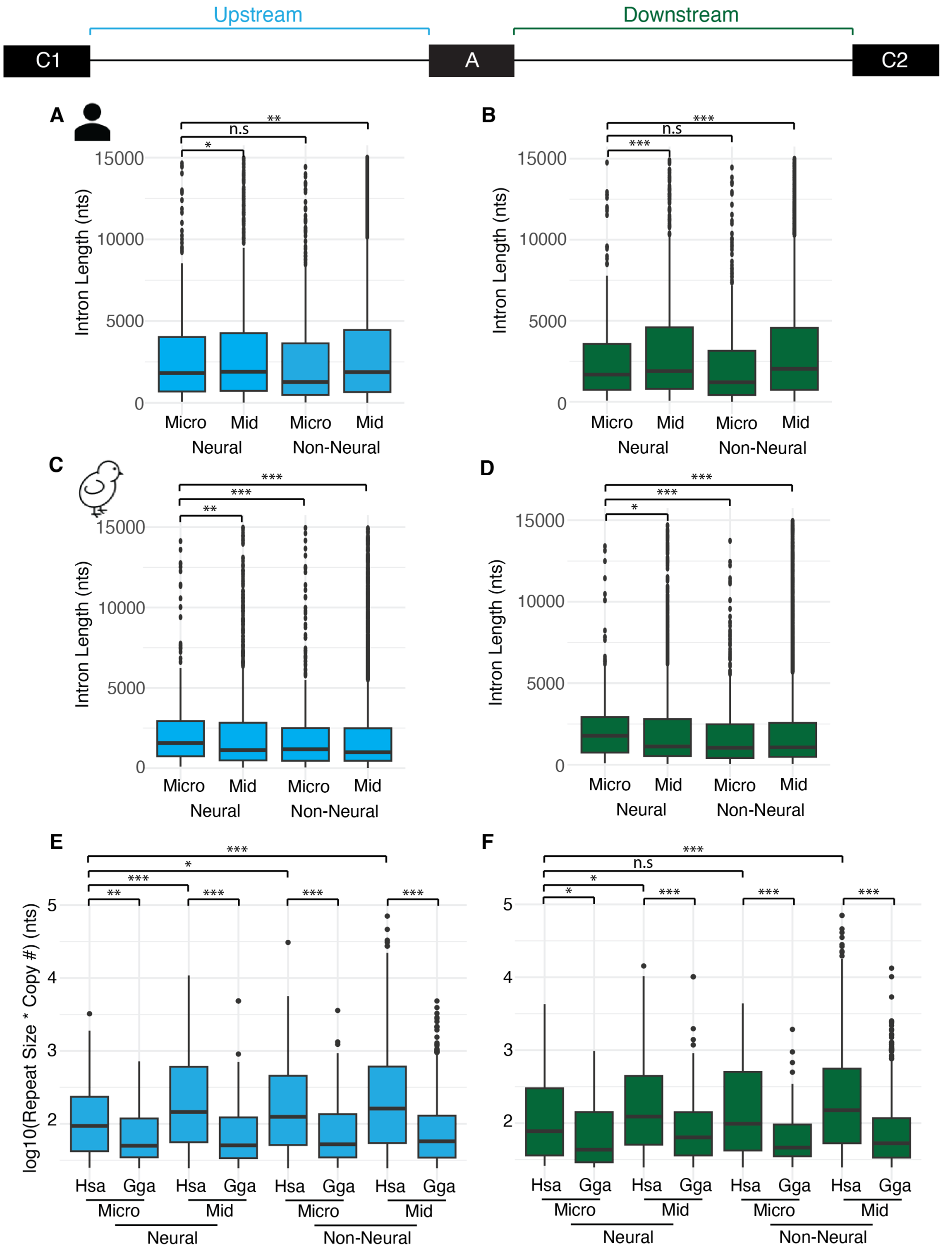


**Figure S10. Short flanking intron length in human neural microexons is maintained despite expanded repetitive elements.** (A) Lengths of upstream introns (blue) and (B) downstream introns (green) flanking human alternatively spliced exons. The upstream constitutive exon 1 (C1), alternative exon (A), downstream constitutive exon 2 (C1) and intronic regions are not to scale. (C) Lengths of upstream introns (blue) and (B) downstream introns (green) flanking chicken alternatively spliced exons. (y-axis display limited to 15,000). (E) Number of nucleotides belonging to repetitive sequences in upstream and (F) downstream introns. (Wilcoxon signed-rank test; * = p-value < 0.05, ** = p-value < 0.01, *** = p-value < 0.005, n.s = p-value > 0.05).

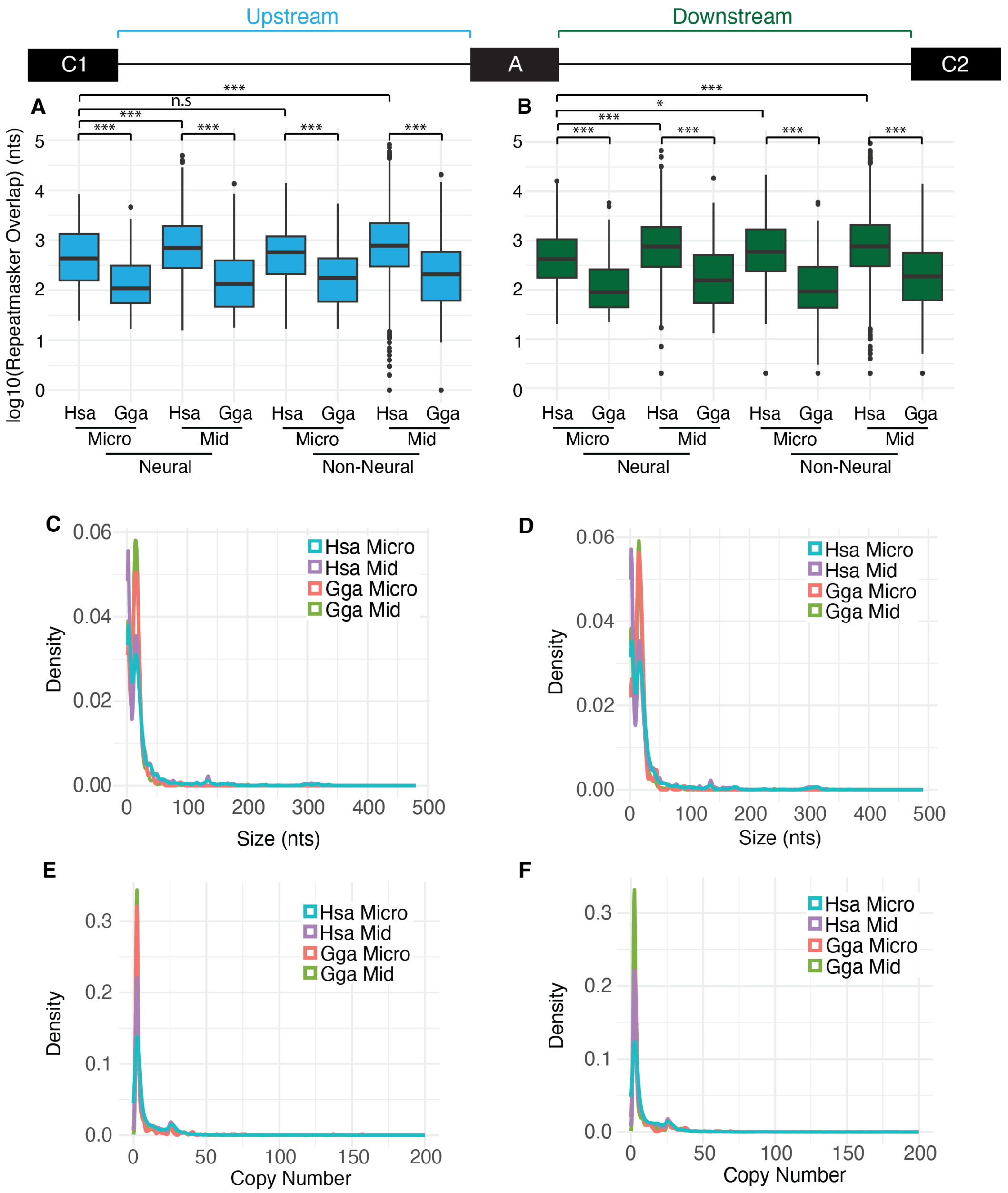


**Figure S11. Humans have higher repetitive sequence content in flanking introns than chickens.** (A) Number of upstream intron nucleotides that overlap with repetitive elements in Repeatmasker annotations. (B) Number of upstream intron nucleotides that overlap with repetitive elements in Repeatmasker annotations. (C) Distribution of repeat sizes of repetitive elements identified in the upstream intron with TRF. (D) Distribution of repeat sizes of repetitive elements identified in the downstream intron with TRF. Plot C and D x-axis limited between 0-500. (E) Distribution of repeat copy numbers of repetitive elements identified in the upstream intron with TRF. (F) Distribution of repeat copy numbers of repetitive elements identified in the downstream intron with TRF. Plot E and F x-axis limited between 0-200; (* = p-value < 0.05, ** = p-value < 0.01, *** = p-value < 0.005, n.s = p-value > 0.05).

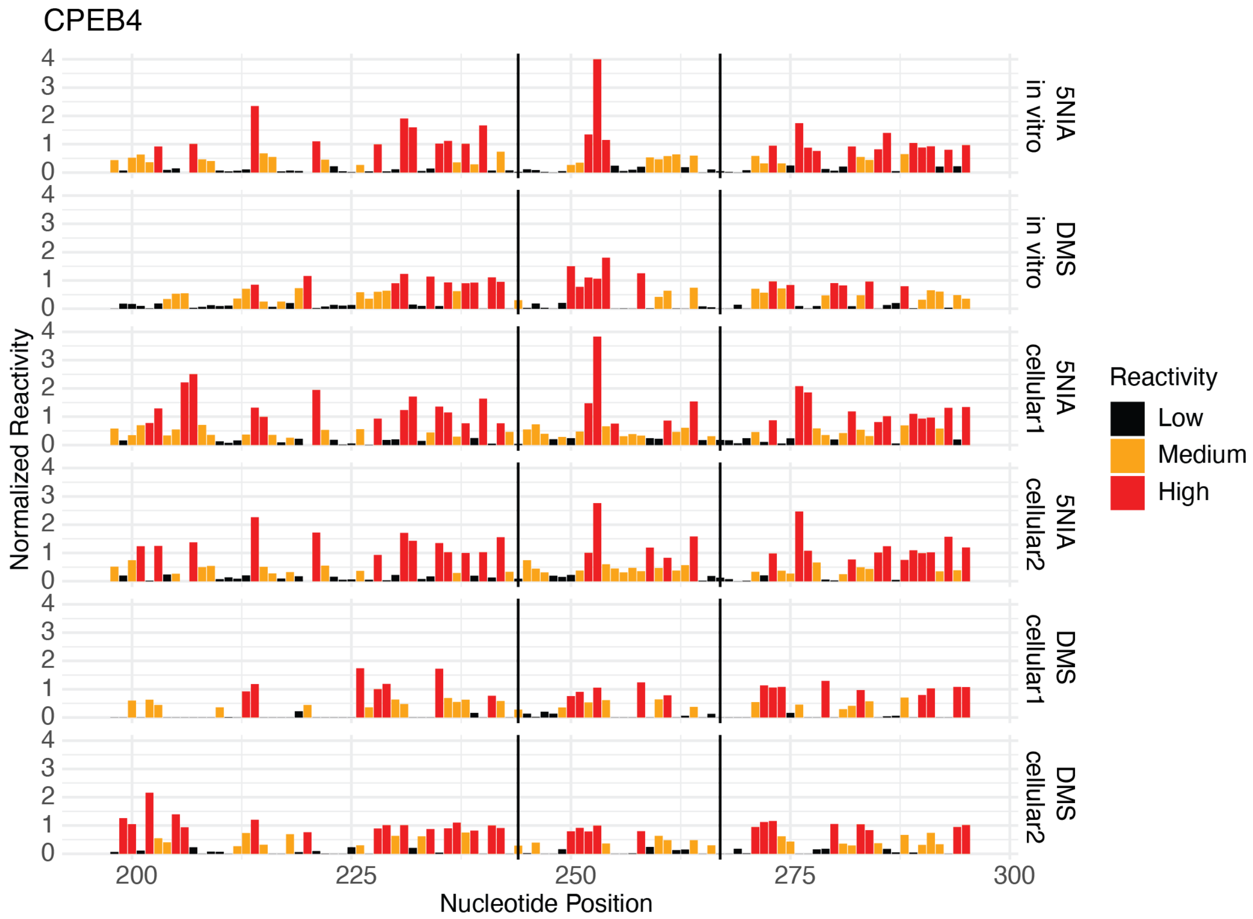


**Figure S12.** ***CPEB4*** **reactivity data for** ***in vitro*** **and cellular SHAPE-MaP and DMS-MaP experiments.**

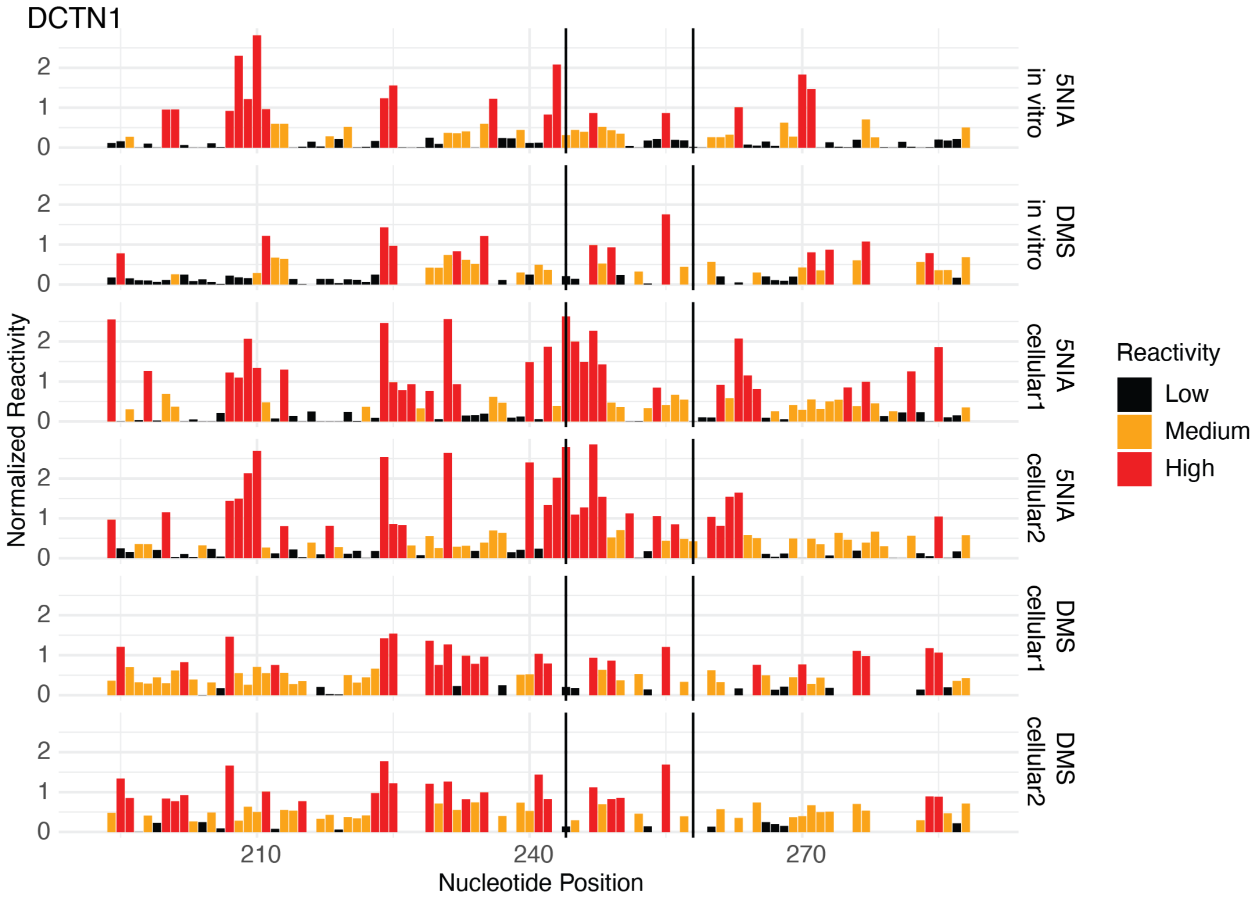


**Figure S13. *DCTN1* reactivity data for *in vitro* and cellular SHAPE-MaP and DMS-MaP experiments.**

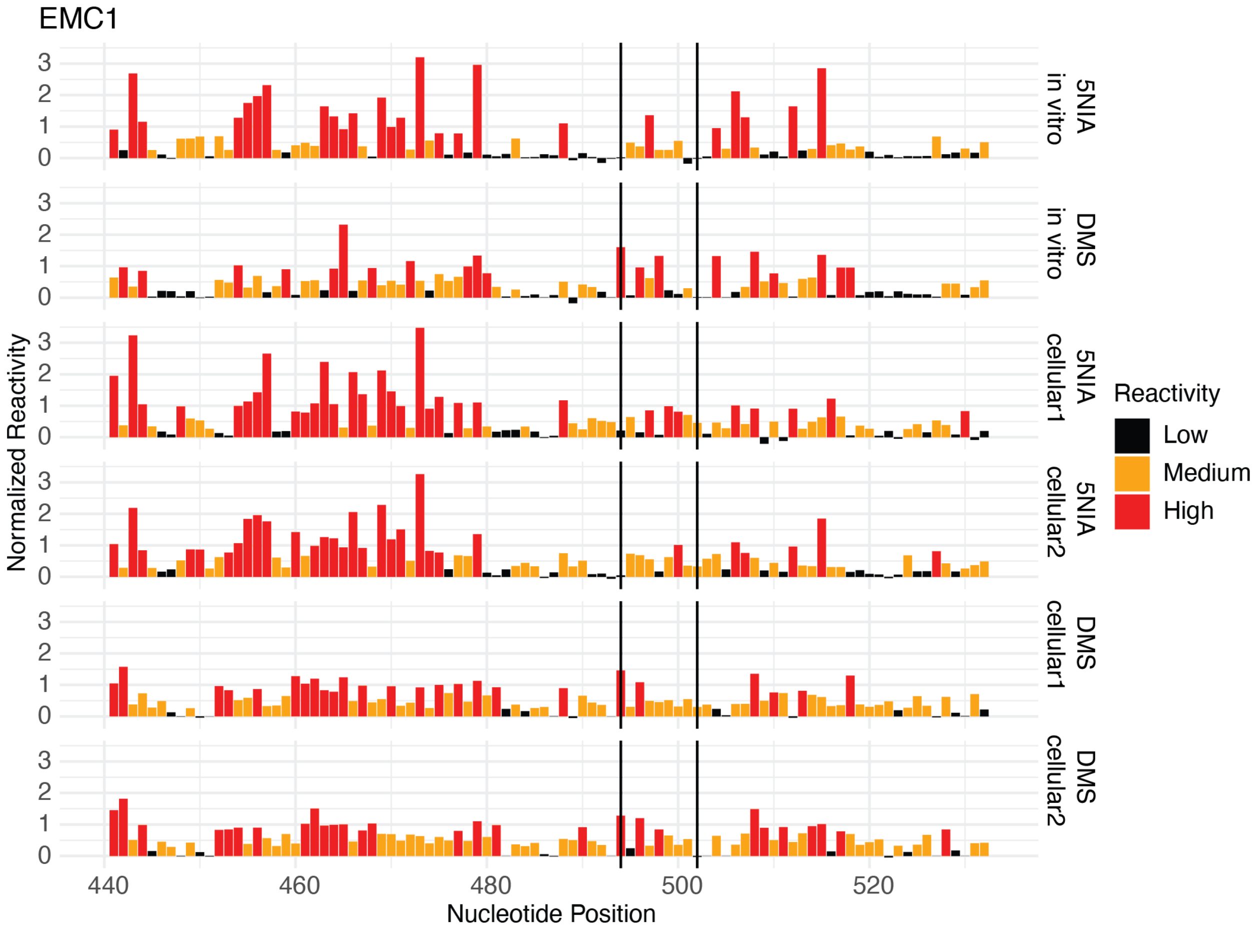


**Figure S14.** ***EMC1*** **reactivity data for** ***in vitro*** **and cellular SHAPE-MaP and DMS-MaP experiments.**

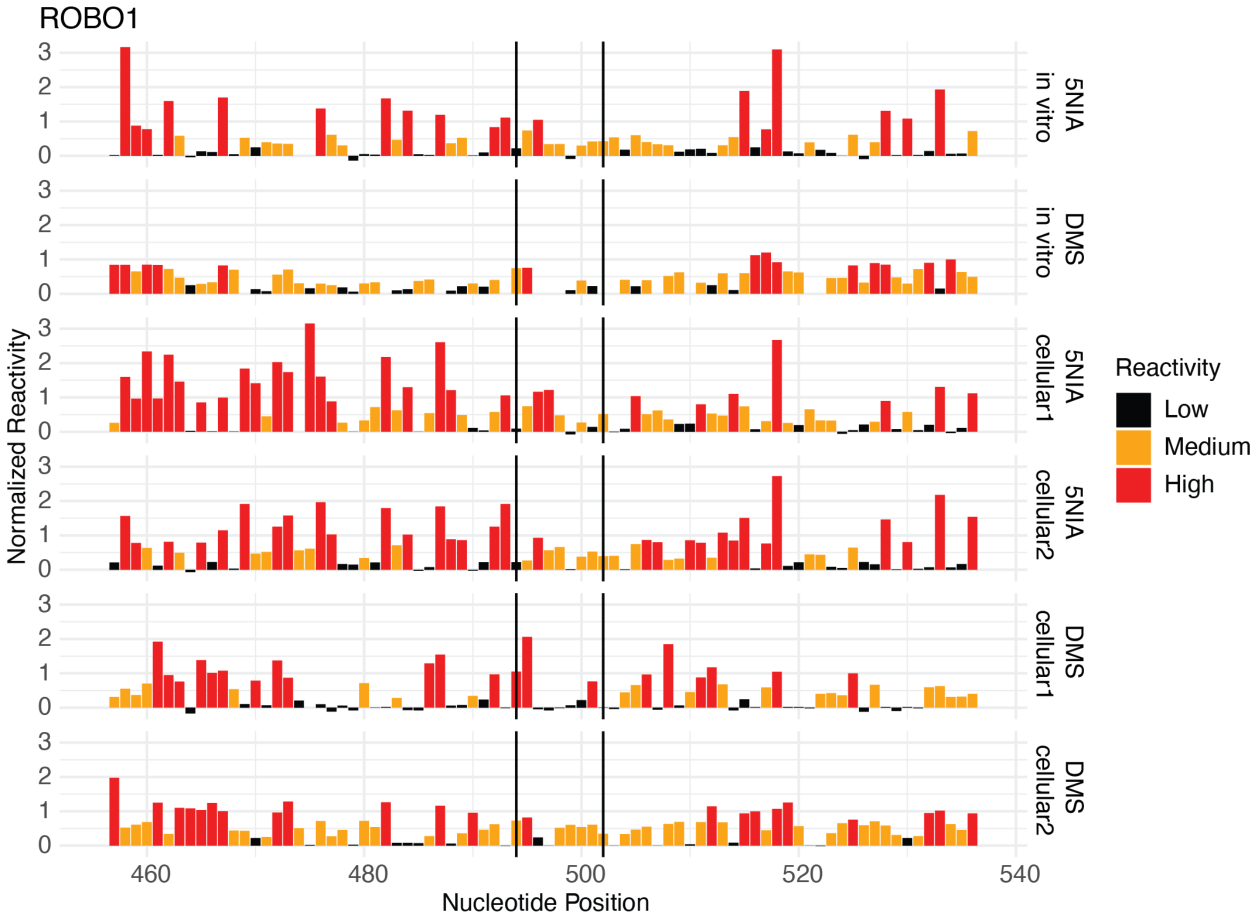


**Figure S15. *ROBO1* reactivity data for *in vitro* and cellular SHAPE-MaP and DMS-MaP experiments.**

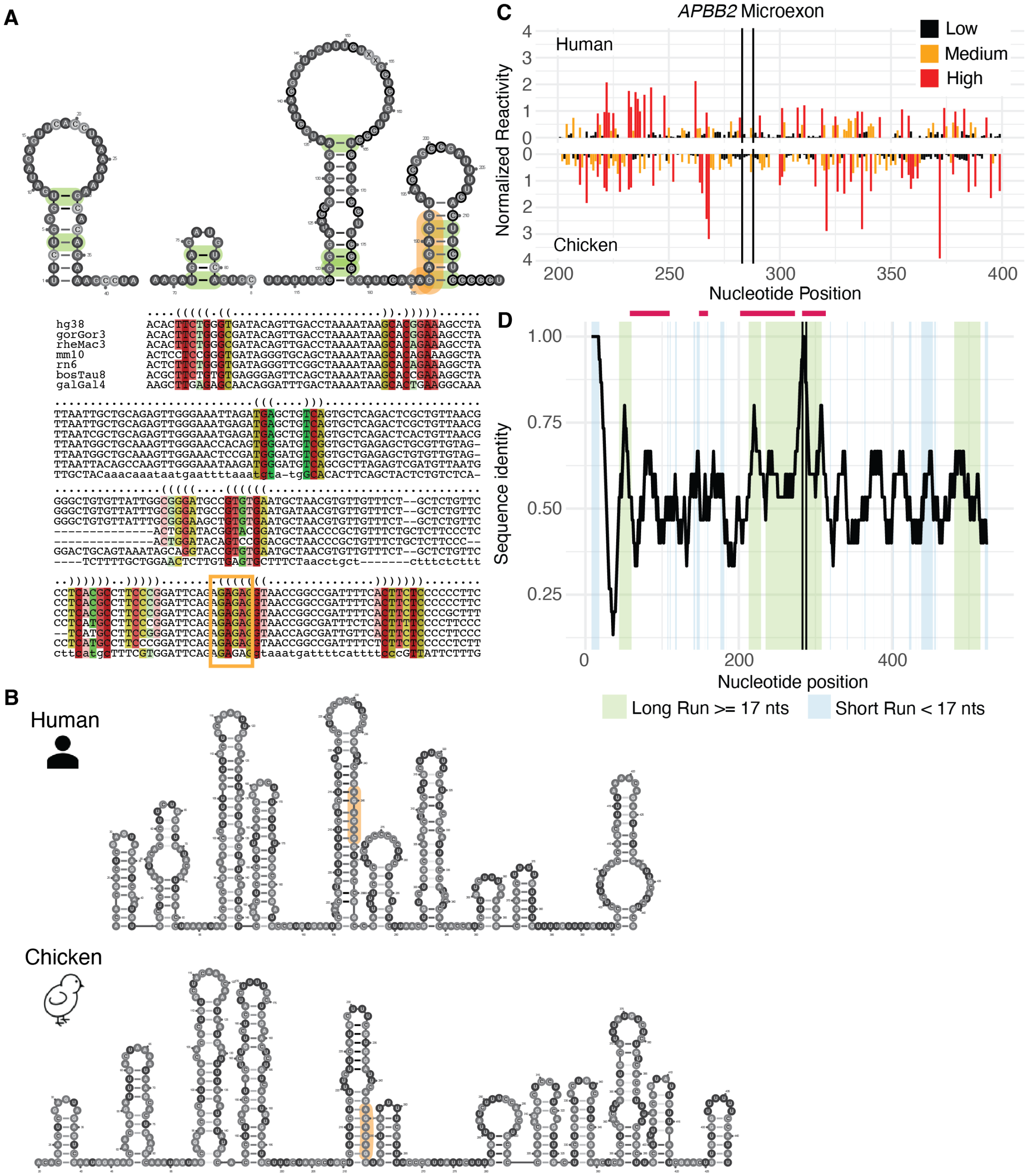


**Figure S16. Reactivity similarity of *APBB2* between human and chicken.** (A) Predicted *APBB2* structure and sequence alignment based on covariation (green) and conservation (red) within common model organisms. (B) Representative structures of human (top) and chicken (bottom) *APBB2*. Microexon highlighted in orange and covariate positions in green if present. (C) Reactivity profiles of human and chicken *APBB2* with splice sites indicated by black vertical lines. (D) *APBB2* sequence identity across aligned human and chicken sequences using a 15-nt sliding window (step size = 1nt). Green or blue shading indicates regions >= 17 nts or < 17 nts, respectively, with Spearman correlations between aligned human and chicken reactivities of >= 0.2; Red bars indicate position of structures predicted with covariation and conservation data. APBB2 = amyloid beta precursor protein binding family B member 2.

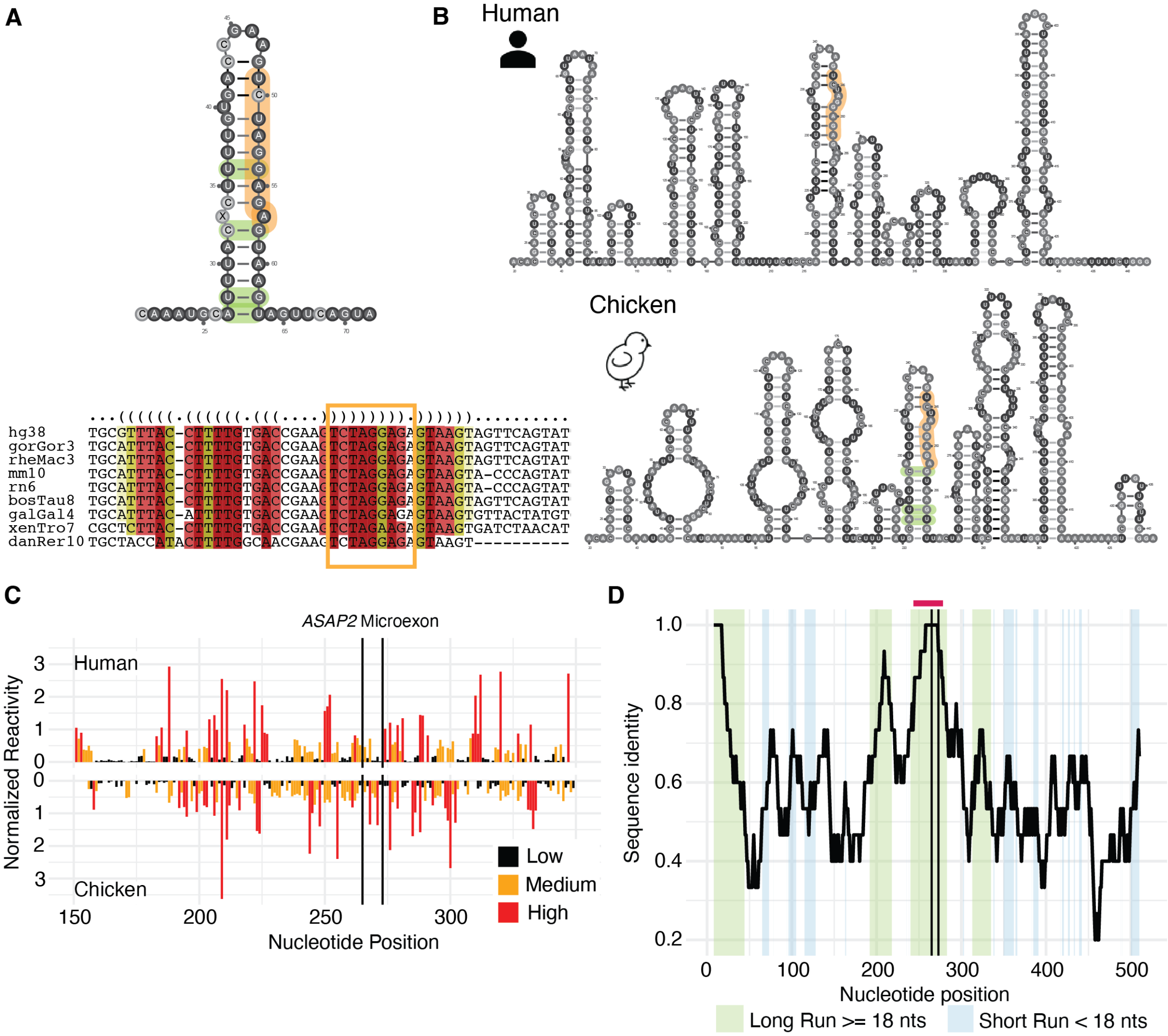


**Figure S17. Reactivity similarity of *ASAP2* between human and chicken.** (A) Predicted *ASAP2* structure and sequence alignment based on covariation (green) and conservation (red) within common model organisms. (B) Representative structures of human (top) and chicken (bottom) *ASAP2*. Microexon highlighted in orange and covariate positions in green if present. (C) Reactivity profiles of human and chicken *ASAP2* with splice sites indicated by black vertical lines. (D) *ASAP2* sequence identity across aligned human and chicken sequences using a 15-nt sliding window (step size = 1nt). Green or blue shading indicates regions >= 18 nts or < 18 nts, respectively, with Spearman correlations between aligned human and chicken reactivities of >= 0.2; Red bars indicate position of structures predicted with covariation and conservation data. ASAP2 = ArfGAP with SH3 domain, ankyrin repeat and PH domain 2.

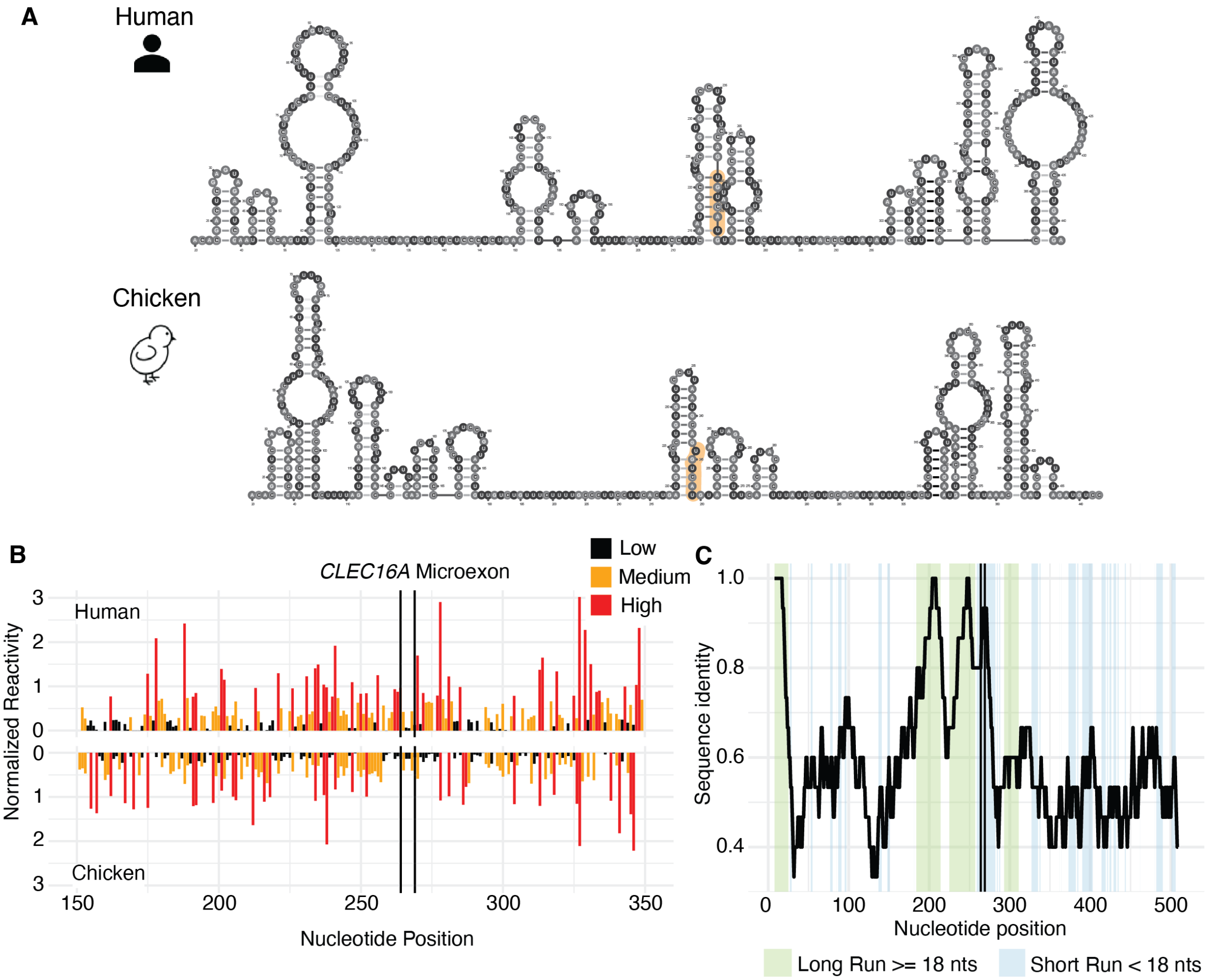


**Figure S18. Reactivity similarity of *CLEC16A* between human and chicken.** (A) Representative structures of human (top) and chicken (bottom) *CLEC16A*. Microexon highlighted in orange. (B) Reactivity profiles of human and chicken *CLEC16A* with splice sites indicated by black vertical lines. (C) *CLEC16A* sequence identity across aligned human and chicken sequences using a 15-nt sliding window (step size = 1nt). Green or blue shading indicates regions >= 18 nts or < 18 nts, respectively, with Spearman correlations between aligned human and chicken reactivities of >= 0.2; Red bars indicate position of structures predicted with covariation and conservation data. CLEC16A = C-type lectin domain family 16 member A.

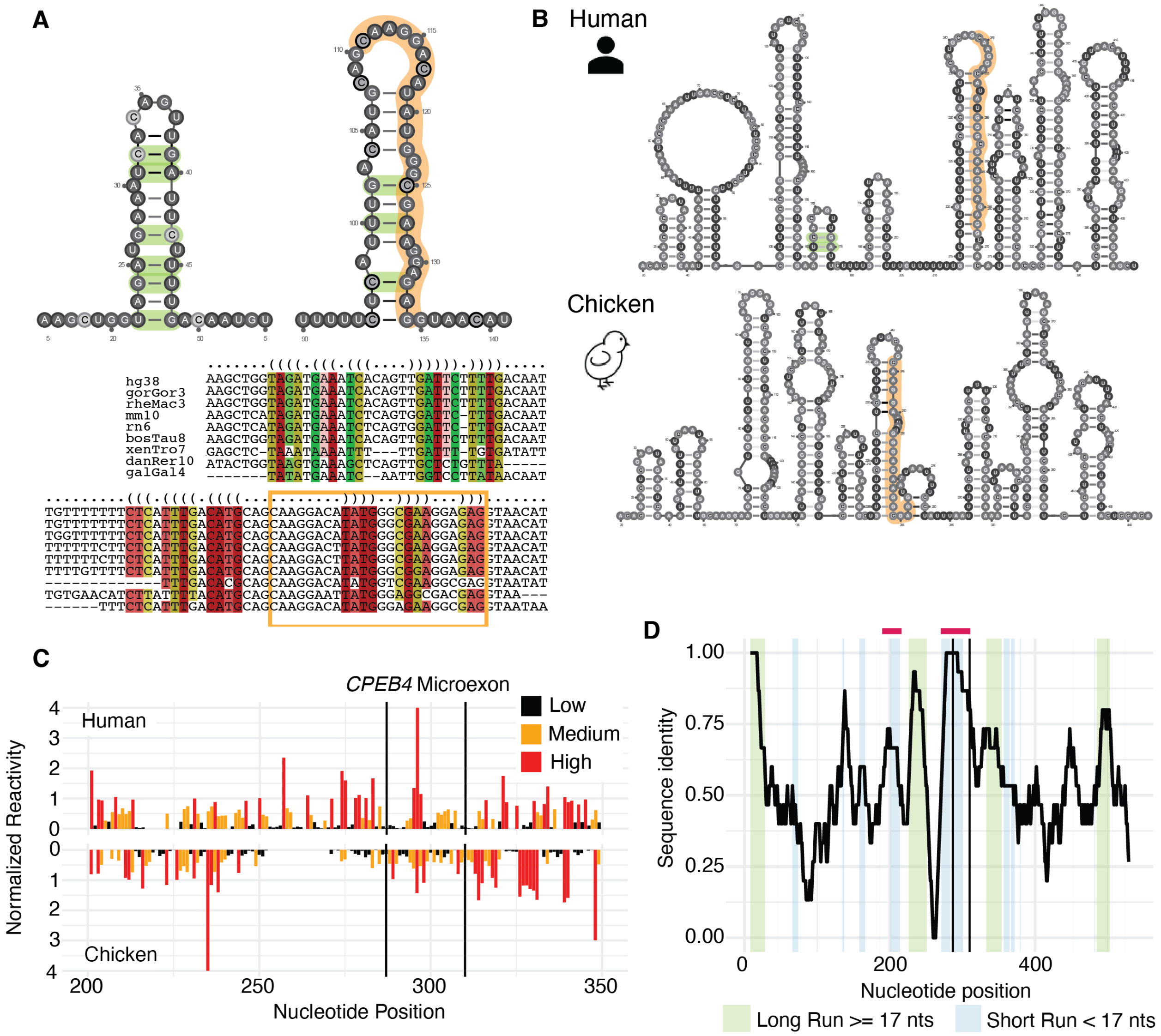


**Figure S19. Reactivity similarity of *CPEB4* between human and chicken.** (A) *CPEB4* structure and sequence alignment based on covariation (green) and conservation (red) within common model organisms. (B) Representative structures of human (top) and chicken (bottom) *CPEB4*. Microexon highlighted in orange and covariate positions in green if present. (C) Reactivity profiles of human and chicken *CPEB4* with splice sites indicated by black vertical lines. (D) *CPEB4* sequence identity across aligned human and chicken sequences using a 15-nt sliding window (step size = 1nt). Green or blue shading indicates regions >= 17 nts or < 17 nts, respectively, with Spearman correlations between aligned human and chicken reactivities of >= 0.2; Red bars indicate position of structures predicted with covariation and conservation data. CPEB4 = cytoplasmic polyadenylation element binding protein 4.

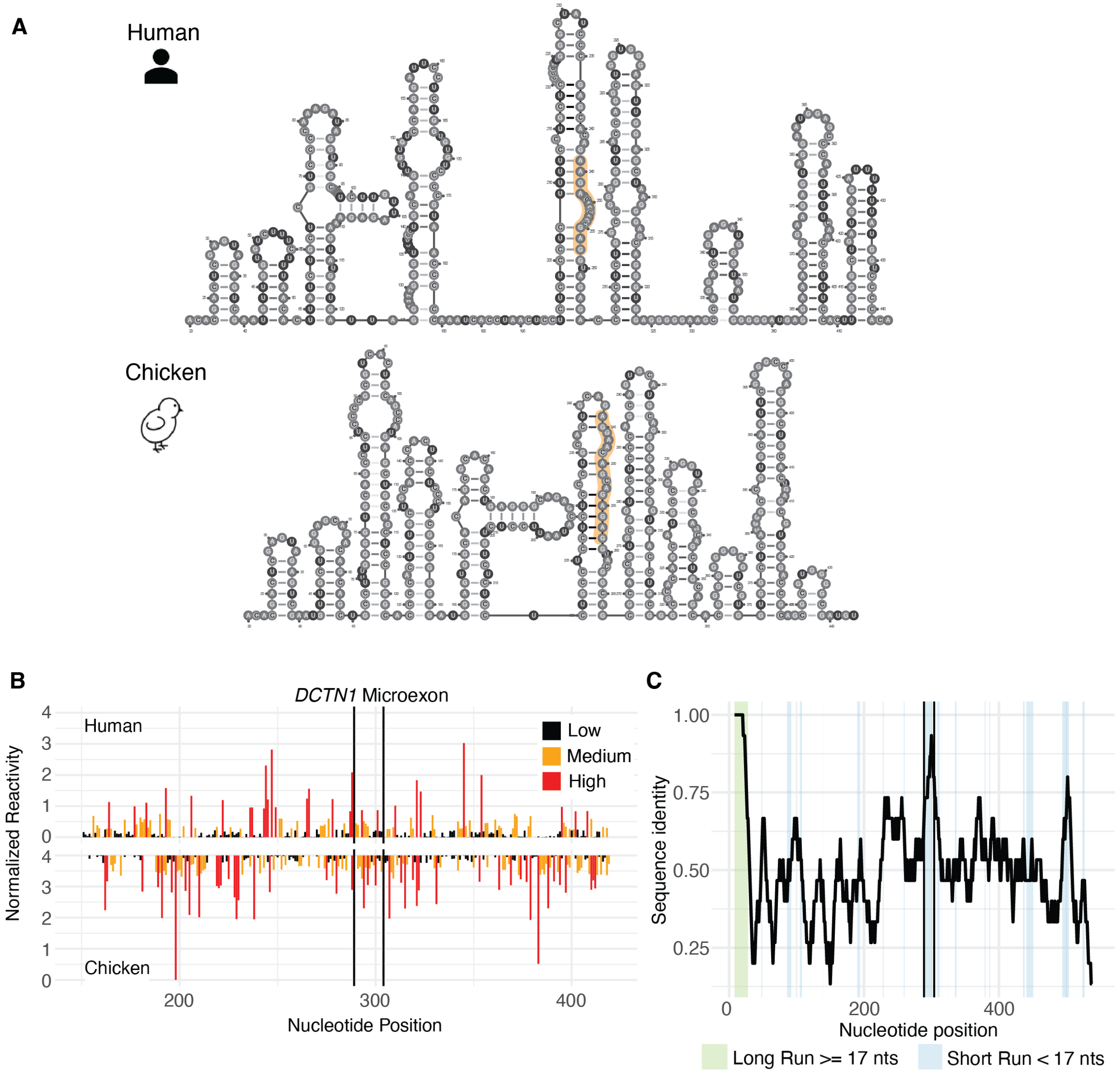


**Figure S20. Reactivity similarity of *DCTN1* between human and chicken.** (A) Representative structures of human (top) and chicken (bottom) *DCTN1*. Microexon highlighted in orange. (B) Reactivity profiles of human and chicken *DCTN1* with splice sites indicated by black vertical lines. (C) *DCTN1* sequence identity across aligned human and chicken sequences using a 15-nt sliding window (step size = 1nt). Green or blue shading indicates regions >= 17 nts or < 17 nts, respectively, with Spearman correlations between aligned human and chicken reactivities of >= 0.2. DCTN1 = dynactin subunit 1.

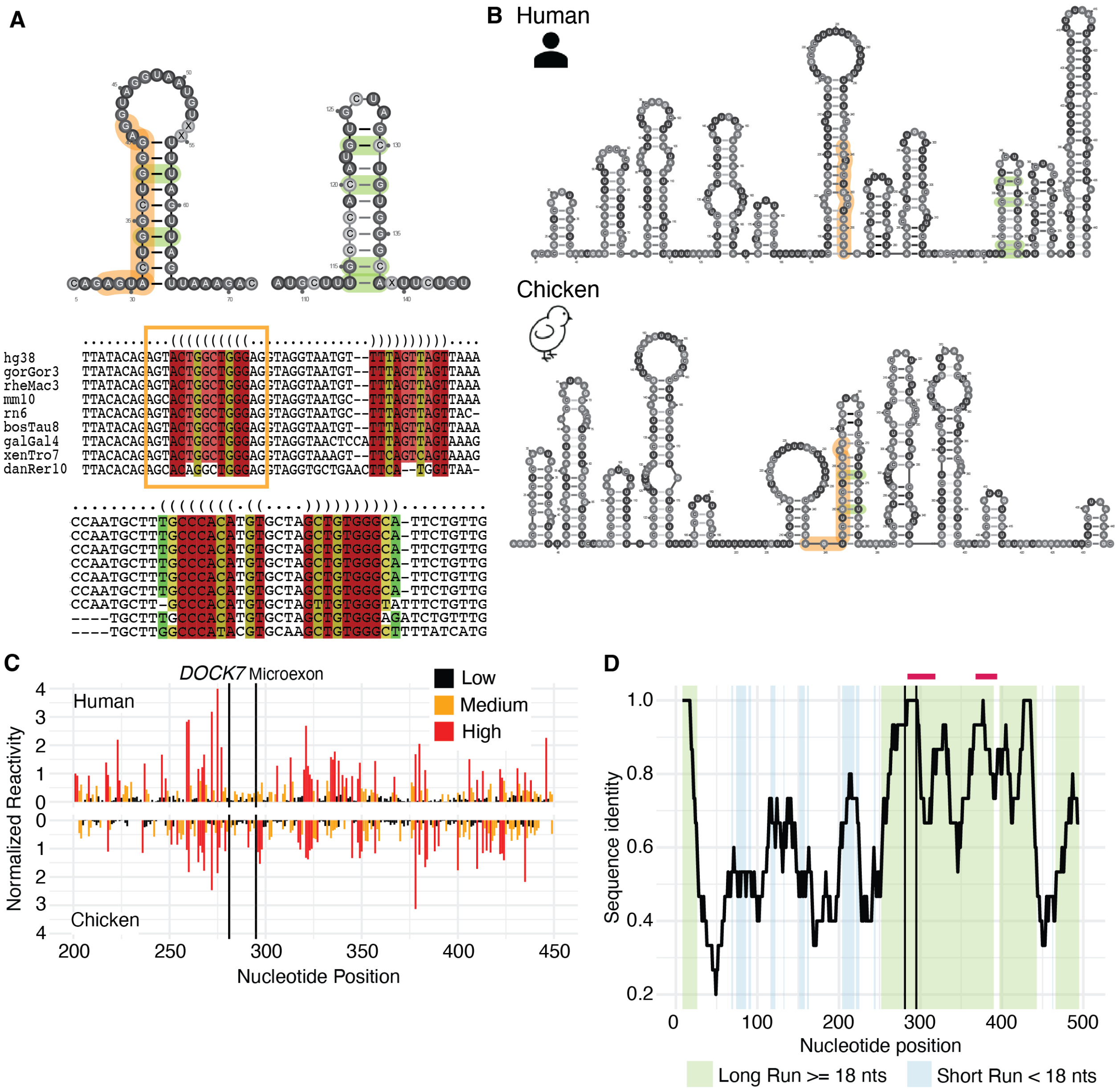


**Figure S21. Reactivity similarity of *DOCK7* between human and chicken.** (A) *DOCK7* structure and sequence alignment based on covariation (green) and conservation (red) within common model organisms. (B) Representative structures of human (top) and chicken (bottom) *DOCK7*. Microexon highlighted in orange and covariate positions in green if present. (C) Reactivity profiles of human and chicken *DOCK7* with splice sites indicated by black vertical lines. (D) *DOCK7* sequence identity across aligned human and chicken sequences using a 15-nt sliding window (step size = 1nt). Green or blue shading indicates regions >= 18 nts or < 18 nts, respectively, with Spearman correlations between aligned human and chicken reactivities of >= 0.2; Red bars indicate position of structures predicted with covariation and conservation data. DOCK7 = dedicator of cytokinesis 7.

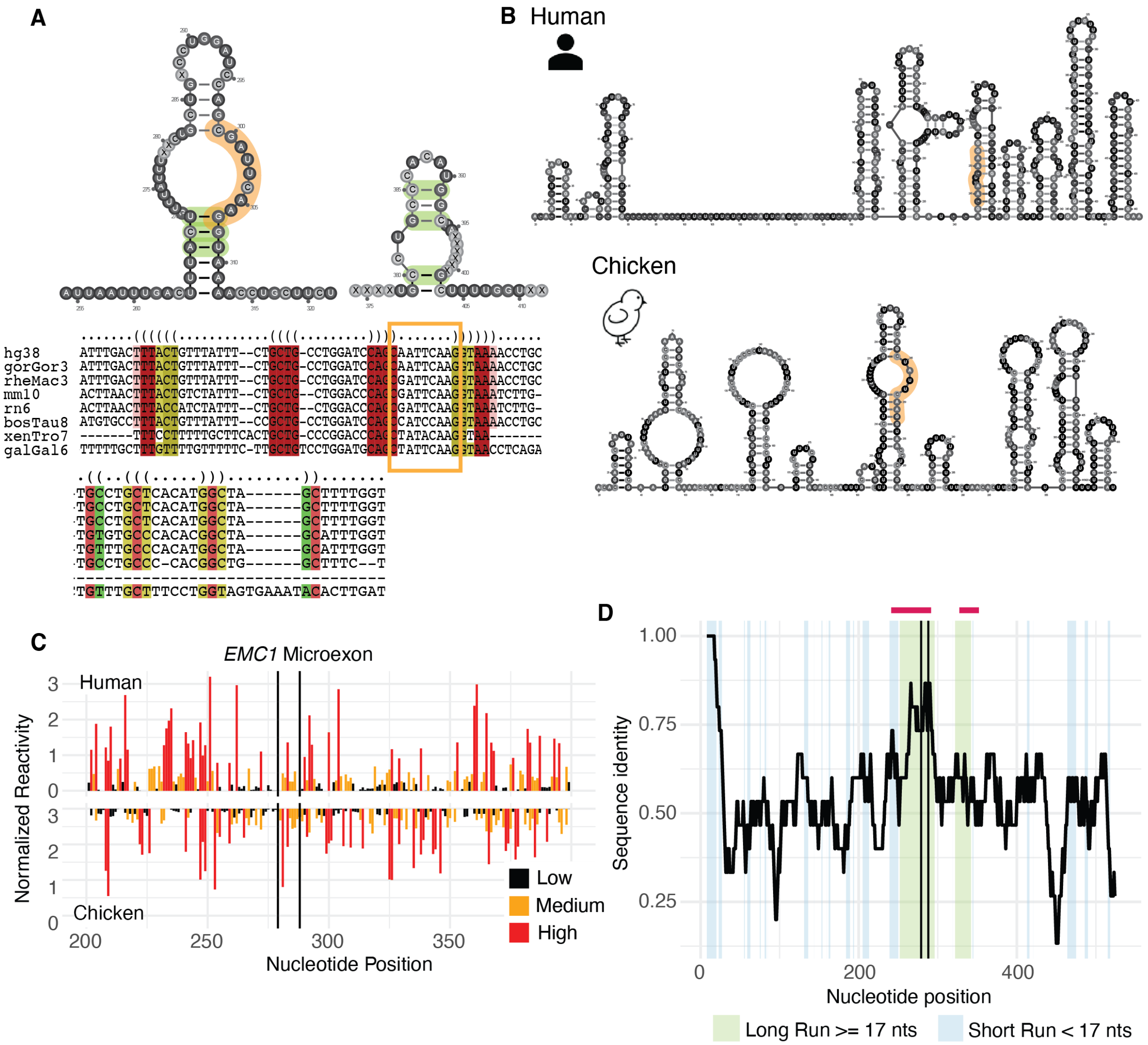


**Figure S22. Reactivity similarity of *EMC1* between human and chicken.** (A) *EMC1* structure and sequence alignment based on covariation (green) and conservation (red) within common model organisms. (B) Representative structures of human (top) and chicken (bottom) *EMC1*. Microexon highlighted in orange and covariate positions in green if present. (C) Reactivity profiles of human and chicken *EMC1* with splice sites indicated by black vertical lines. (D) *EMC1* sequence identity across aligned human and chicken sequences using a 15-nt sliding window (step size = 1nt). Green or blue shading indicates regions >= 17 nts or < 17 nts, respectively, with Spearman correlations between aligned human and chicken reactivities of >= 0.2; Red bars indicate position of structures predicted with covariation and conservation data. EMC1 = ER membrane protein complex subunit 1.

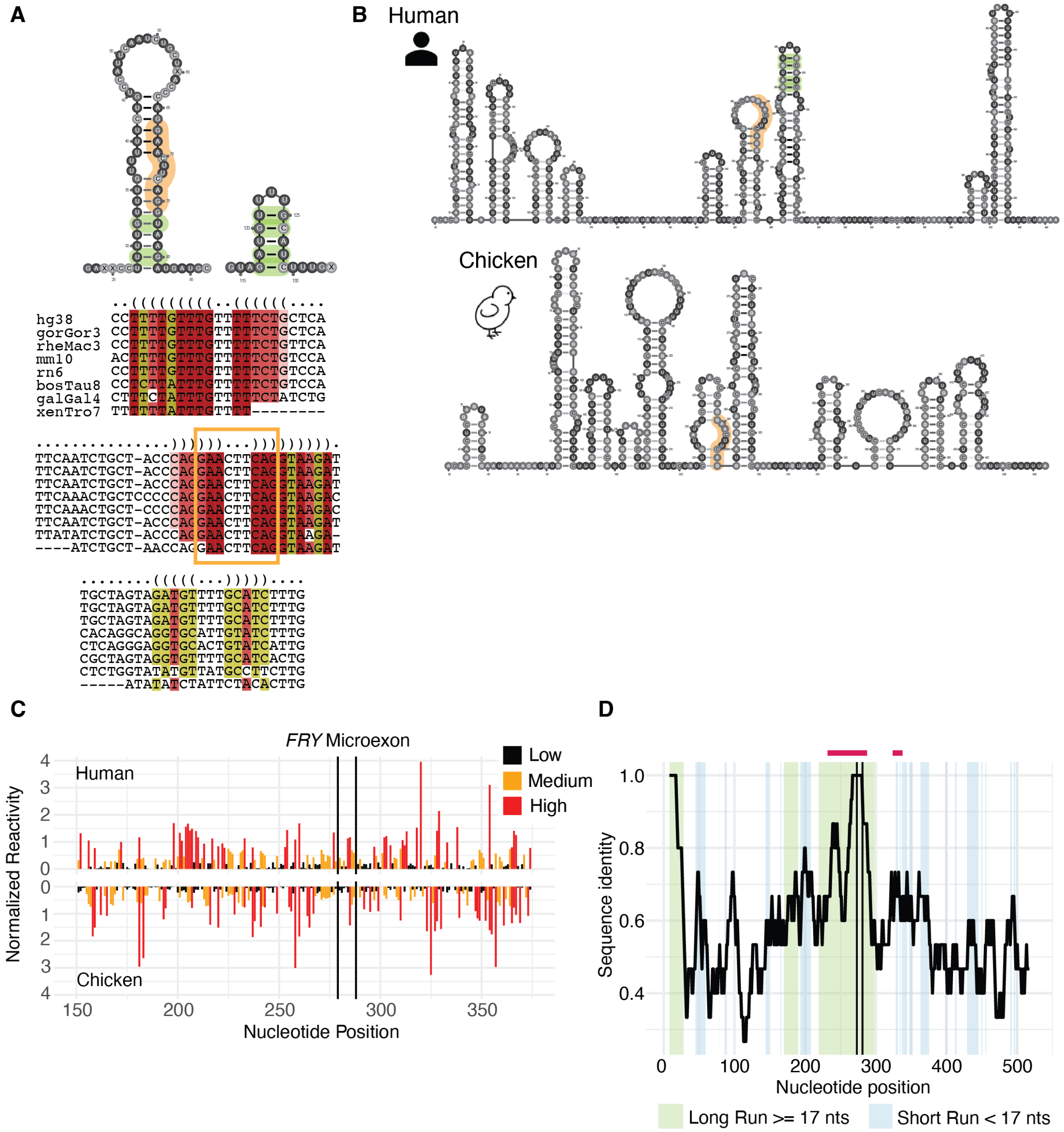


**Figure S23. Reactivity similarity of *FRY* between human and chicken.** (A) *FRY* structure and sequence alignment based on covariation (green) and conservation (red) within common model organisms. (B) Representative structures of human (top) and chicken (bottom) *FRY*. Microexon highlighted in orange and covariate positions in green if present. (C) Reactivity profiles of human and chicken *FRY* with splice sites indicated by black vertical lines. (D) *FRY* sequence identity across aligned human and chicken sequences using a 15-nt sliding window (step size = 1nt). Green or blue shading indicates regions >= 17 nts or < 17 nts, respectively, with Spearman correlations between aligned human and chicken reactivities of >= 0.2; Red bars indicate position of structures predicted with covariation and conservation data. FRY = FRY microtubule binding protein.

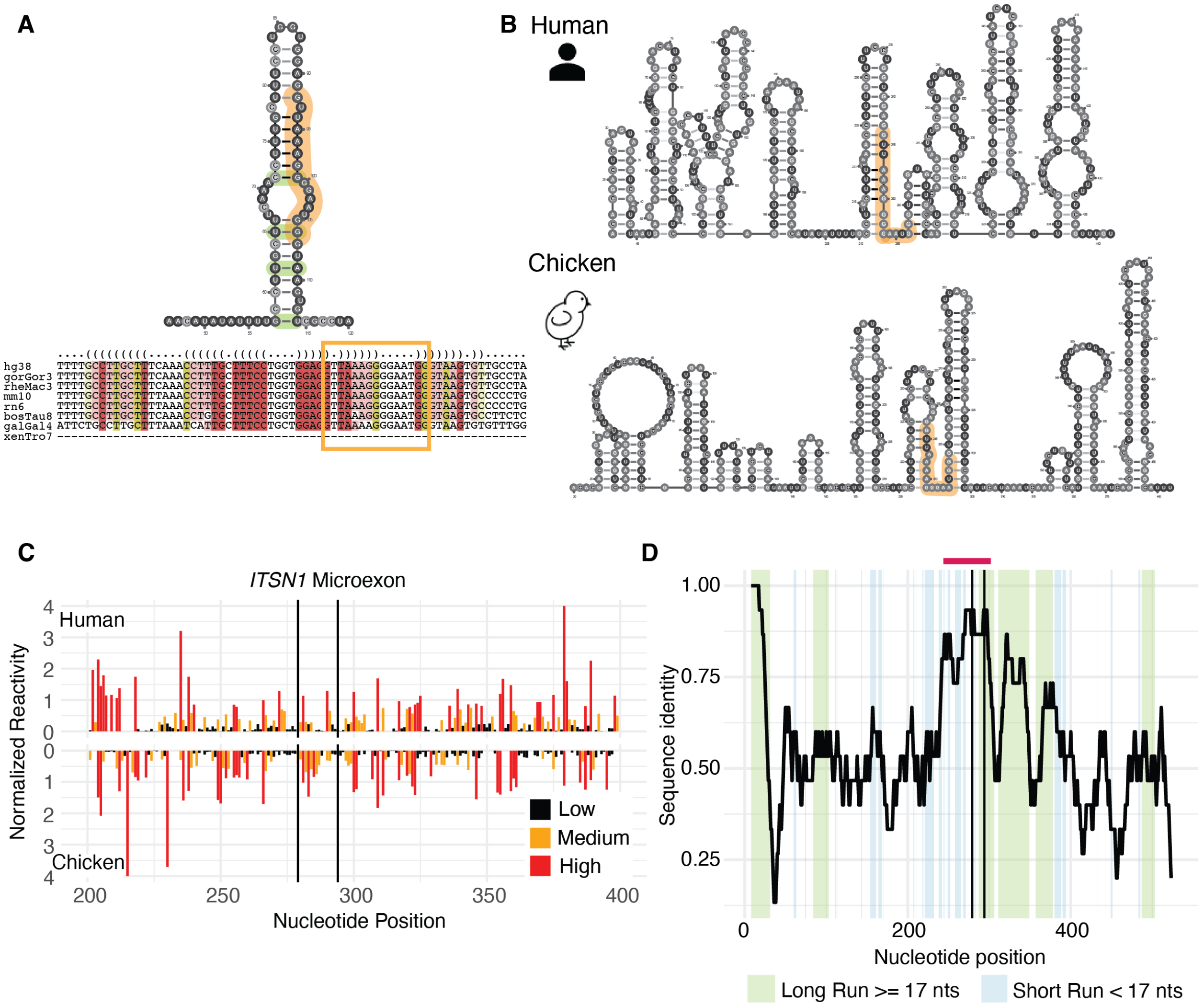


**Figure S24. Reactivity similarity of *ITSN1* between human and chicken.** (A) *ITSN1* structure and sequence alignment based on covariation (green) and conservation (red) within common model organisms. (B) Representative structures of human (top) and chicken (bottom) *ITSN1*. Microexon highlighted in orange and covariate positions in green if present. (C) Reactivity profiles of human and chicken *ITSN1* with splice sites indicated by black vertical lines. (D) *ITSN1* sequence identity across aligned human and chicken sequences using a 15-nt sliding window (step size = 1nt). Green or blue shading indicates regions >= 17 nts or < 17 nts, respectively, with Spearman correlations between aligned human and chicken reactivities of >= 0.2; Red bars indicate position of structures predicted with covariation and conservation data. ITSN1 = intersectin 1.

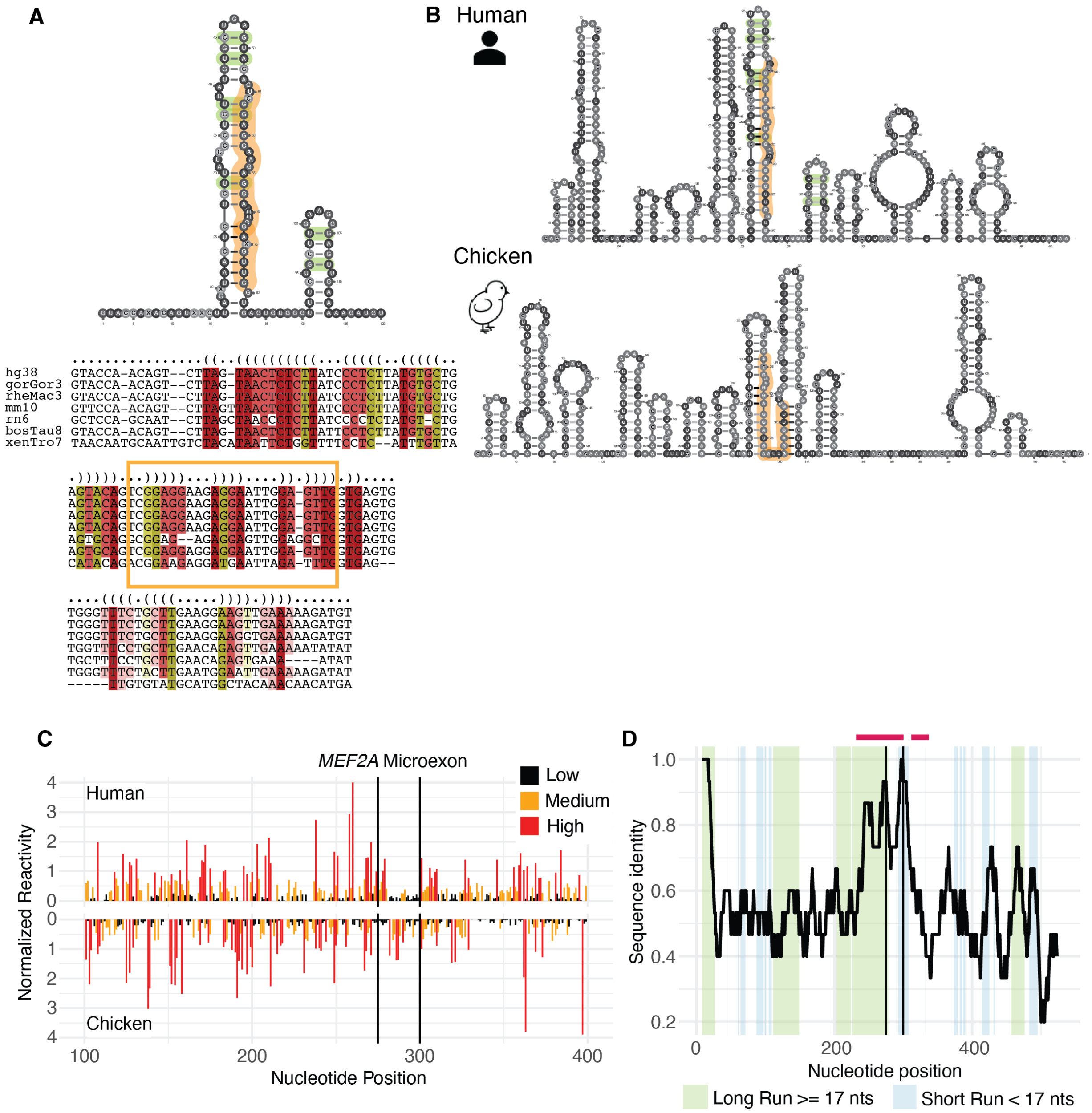


**Figure S25. Reactivity similarity of *MEF2A* between human and chicken.** (A) *MEF2A* structure and sequence alignment based on covariation (green) and conservation (red) within common model organisms. (B) Representative structures of human (top) and chicken (bottom) *MEF2A*. Microexon highlighted in orange and covariate positions in green if present. (C) Reactivity profiles of human and chicken *MEF2A* with splice sites indicated by black vertical lines. (D) *MEF2A* sequence identity across aligned human and chicken sequences using a 15-nt sliding window (step size = 1nt). Green or blue shading indicates regions >= 17 nts or < 17 nts, respectively, with Spearman correlations between aligned human and chicken reactivities of >= 0.2; Red bars indicate position of structures predicted with covariation and conservation data. MEF2A = myocyte enhancer factor 2A.

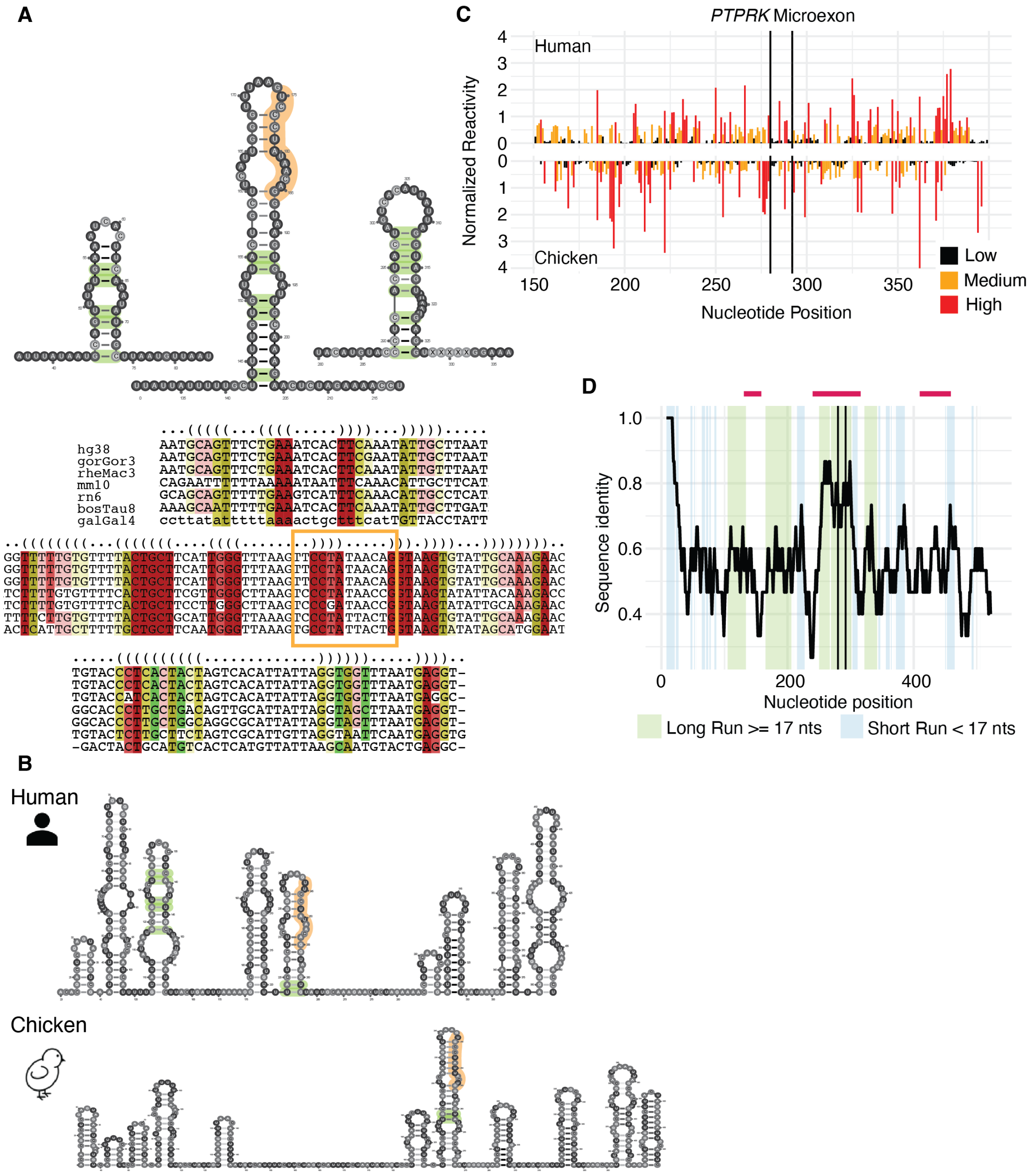


**Figure S26. Reactivity similarity of *PTPRK* between human and chicken.** (A) *PTPRK* structure and sequence alignment based on covariation (green) and conservation (red) within common model organisms. (B) Representative structures of human (top) and chicken (bottom) *PTPRK*. Microexon highlighted in orange and covariate positions in green if present. (C) Reactivity profiles of human and chicken *PTPRK* with splice sites indicated by black vertical lines. (D) *PTPRK* sequence identity across aligned human and chicken sequences using a 15-nt sliding window (step size = 1nt). Green or blue shading indicates regions >= 17 nts or < 17 nts, respectively, with Spearman correlations between aligned human and chicken reactivities of >= 0.2; Red bars indicate position of structures predicted with covariation and conservation data. PTPRK = protein tyrosine phosphatase, receptor type K.

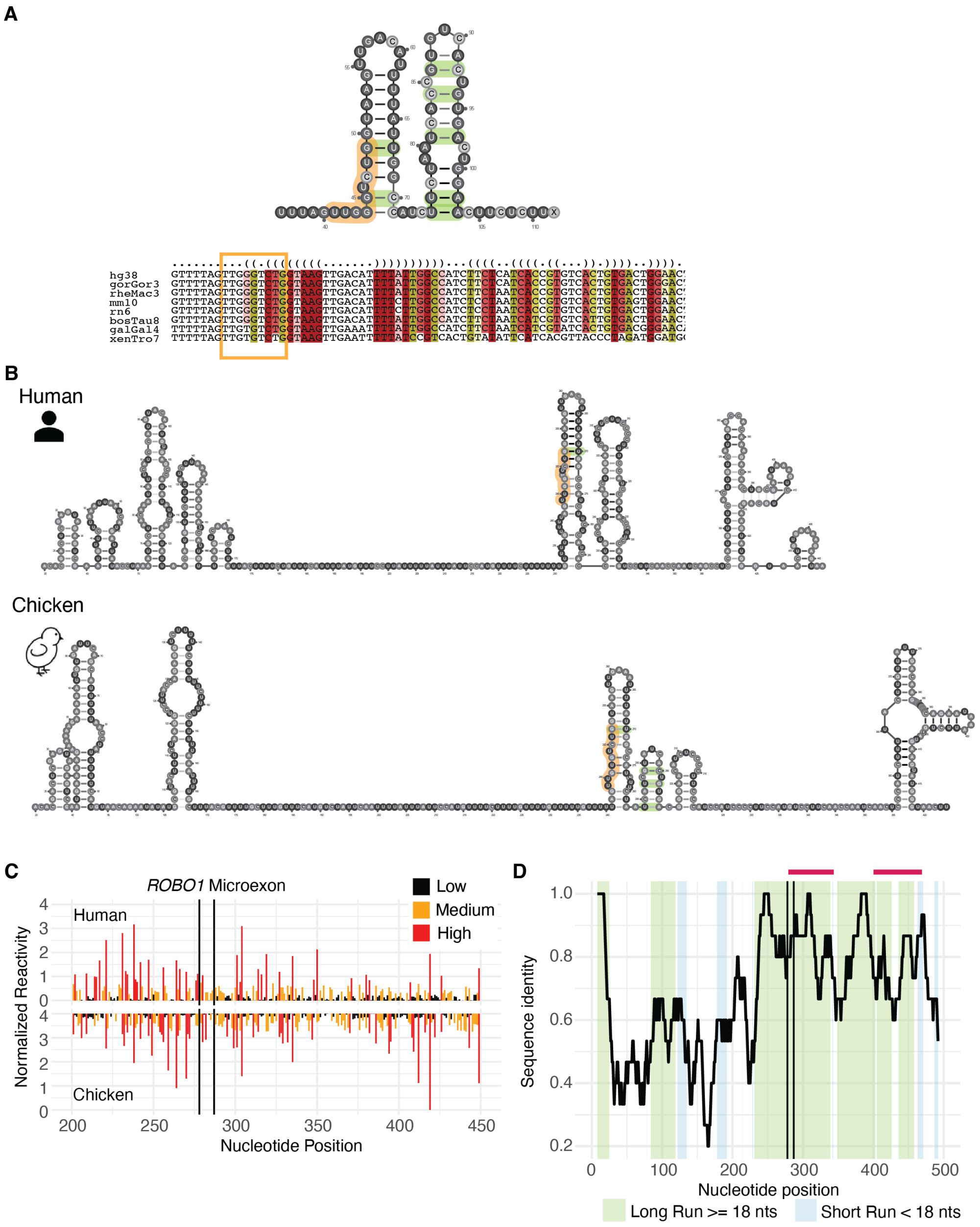


**Figure S27. Reactivity similarity of *ROBO1* between human and chicken.** (A) *ROBO1* structure and sequence alignment based on covariation (green) and conservation (red) within common model organisms. (B) Representative structures of human (top) and chicken (bottom) *ROBO1*. Microexon highlighted in orange and covariate positions in green if present. (C) Reactivity profiles of human and chicken *ROBO1* with splice sites indicated by black vertical lines. (D) *ROBO1* sequence identity across aligned human and chicken sequences using a 15-nt sliding window (step size = 1nt). Green or blue shading indicates regions >= 18 nts or < 18 nts, respectively, with Spearman correlations between aligned human and chicken reactivities of >= 0.2; Red bars indicate position of structures predicted with covariation and conservation data. ROBO1 = roundabout guidance receptor 1.

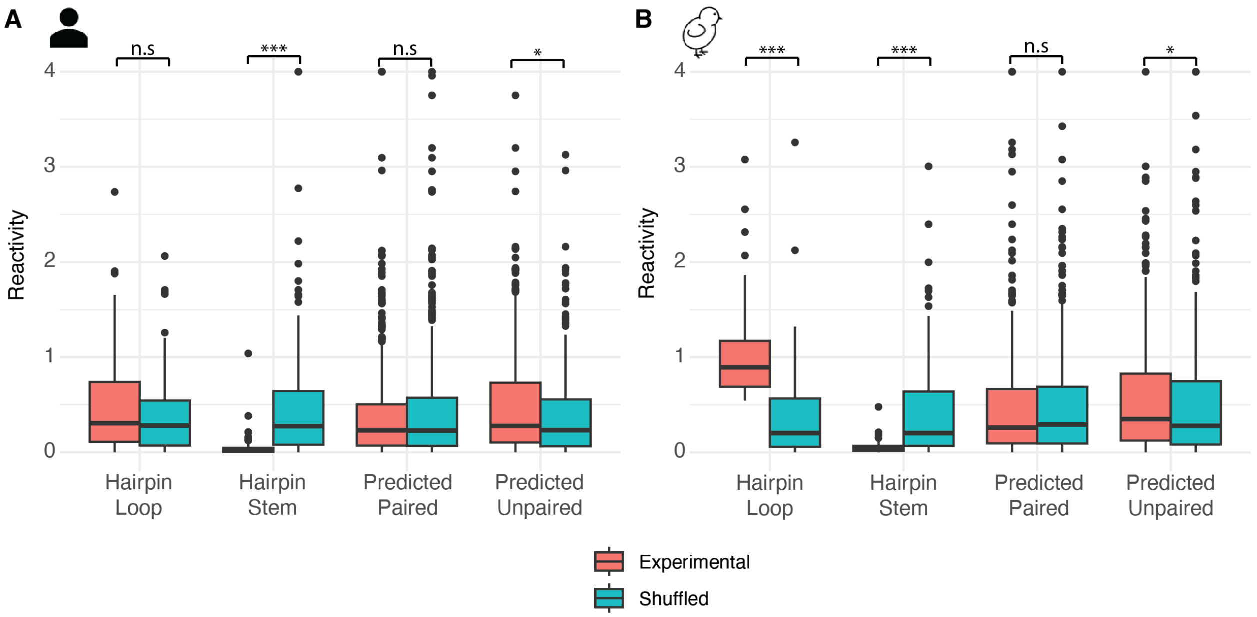


**Figure S28. Structures predicted from covariation and conservation are not supported with experimental data.** (A) Human microexon (n=13) and (B) Chicken microexon (n=13) reactivities at coordinates predicted to be paired or unpaired with RNAalifold (red; see Methods). Reactivities were shuffled as a negative control (blue). The “Hairpin Loop” and “Hairpin Stem” refer to the hairpin stem loop incorporated at the 5’ end of the gene constructs. (* = p-value < 0.05, ** = p-value < 0.01, *** = p-value < 0.005, n.s = p-value > 0.05).

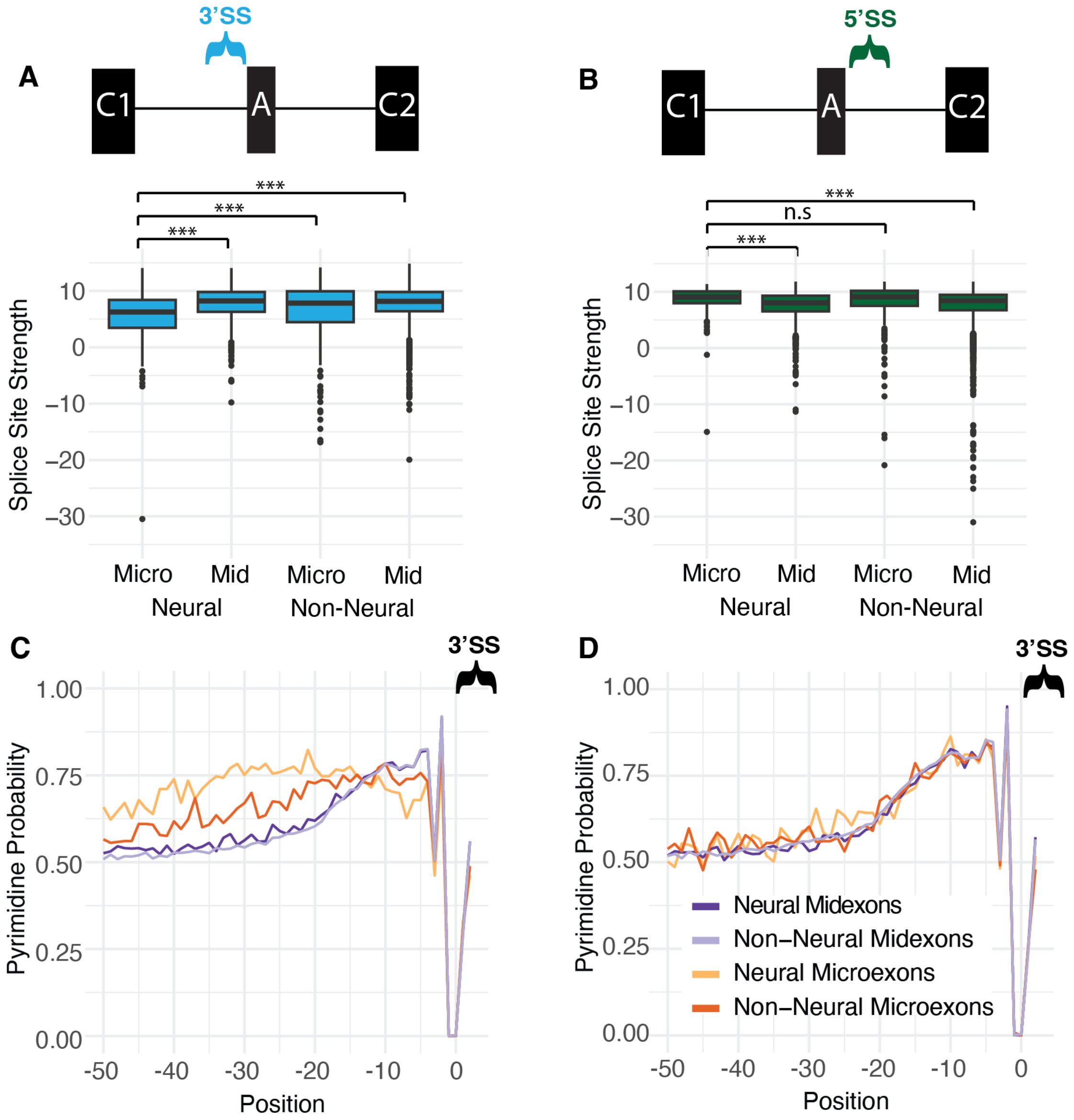


**Figure S29. Chickens have similar weak 3' splice site strength and pyrimidine probability patterns as humans.** (A) 3' splice site strength scores for chicken microexon and midexon sequences. (B) 5' splice site strength scores for chicken microexon and midexon sequences. (C) Probability of cytosine or uracil per nucleotide position upstream of alternatively microexons and midexons. (D) Probability of cytosine or uridine per nucleotide position approaching the 3'ss of the exons downstream of the alternatively spliced microexons and midexons. (Wilcoxon signed-rank test; * = p-value < 0.05, ** = p-value < 0.01, *** = p-value < 0.005).

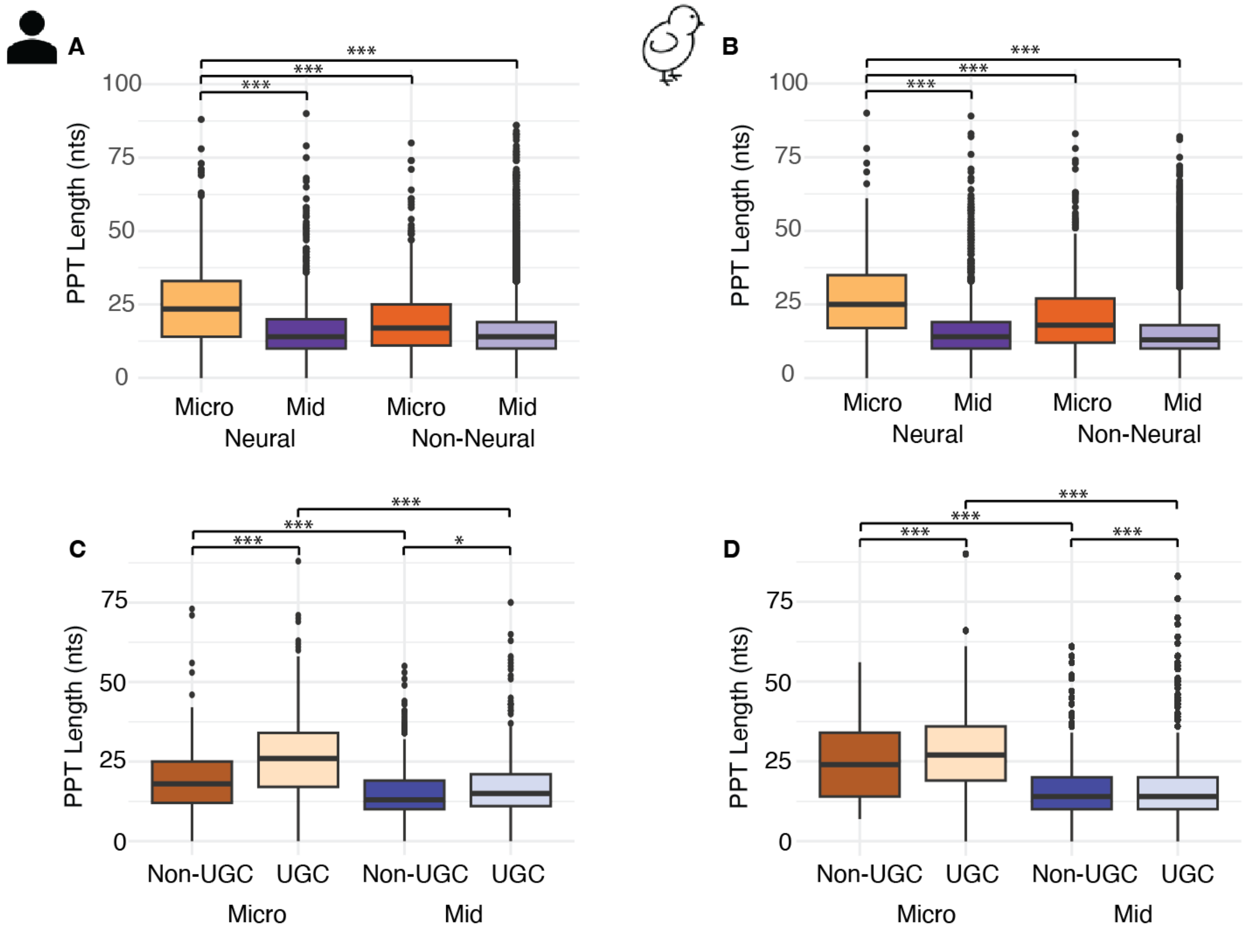


**Figure S30. Neural microexons have an extended polypyrimidine tract in human and chicken.** (A) Polypyrimidine tract length corresponding to top-scoring branchpoint in human and (B) chicken. (C) Polypyrimidine tract length corresponding to top-scoring branchpoint of neural UGC containing and non-UGC containing microexons and midexons in human and (D) chicken. All measurements were generated with SVM-BPfinder; (* = p-value < 0.05, ** = p-value < 0.01, *** = p-value < 0.005, n.s = p-value > 0.05)

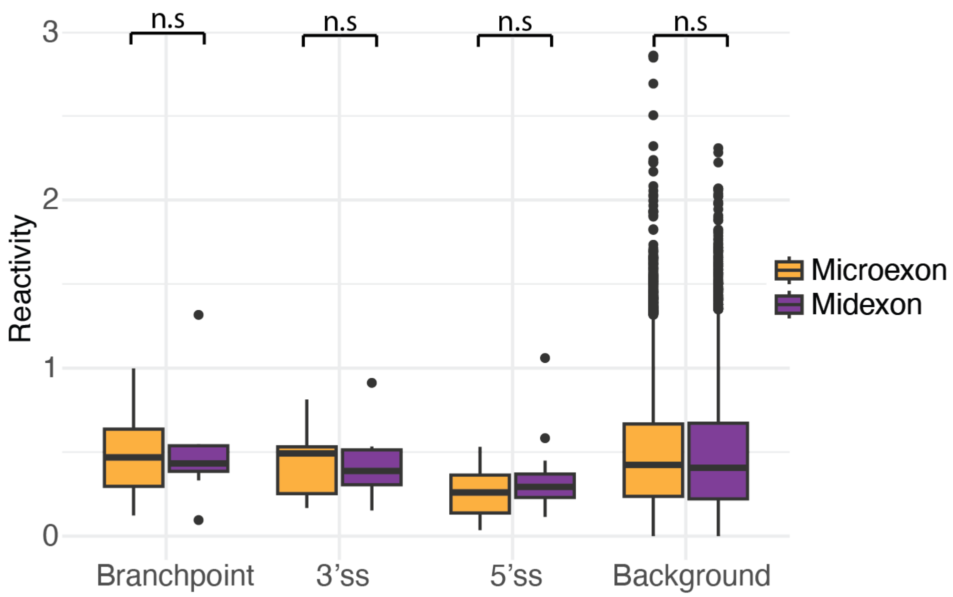


**Figure S31. Splicing regulatory elements do not have significantly different accessibilities in neural microexons compared to neural midexons in humans.** Reactivity values are averaged within 5-nt windows. (* = p-value < 0.05, ** = p-value < 0.01, *** = p-value < 0.005, n.s = p-value > 0.05).

**Supp Table S1. Structure probing replicate correlations (Pearson).**

| *Gene* | *5NIA human* | *5NIA chick* | *DMS human* | *DMS chick* | *Cellular 5NIA* | *Cellular DMS* |
|---|---|---|---|---|---|---|
| *ROBO1* | 0.993 | 0.85 | 0.911 | 0.712 | 0.700 | 0.442 (11/82 AC) |
| *PTPRK* | 0.989 | 0.94 | 0.761 | 0.863 | NA | NA |
| *FRY* | 0.98 | 0.889 | 0.741 | 0.912 | NA | NA |
| *CLEC16A* | 0.976 | 0.976 | 0.786 | 0.888 | NA | NA |
| *APBB2* | 0.992 | 0.926 | 0.975 | 0.856 | NA | NA |
| *ITSN1* | 0.977 | 0.981 | 0.872 | 0.939 | NA | NA |
| *AGAP1* | 0.98 | 0.964 | 0.785 | 0.866 | NA | NA |
| *EMC1* | 0.995 | 0.972 | 0.936 | 0.755 | 0.823 | 0.789 (20/94 AC) |
| *DOCK7* | 0.918 | 0.975 | 0.884 | 0.898 | NA | NA |
| *DCTN1* | 0.914 | 0.933 | 0.956 | 0.968 | 0.785 | 0.788 (27/97 AC) |
| *ASAP2* | 0.96 | 0.962 | 0.955 | 0.889 | NA | NA |
| *MEF2A* | 0.879 | 0.981 | 0.943 | 0.95 | NA | NA |
| *CPEB4* | 0.972 | 0.876 | 0.898 | 0.932 | 0.864 | 0.484 (19/100 AC) |

**Supp Table S2. 5NIA and DMS reactivity correlation and A/C content (Spearman).**

| *Gene* | *Human correlation (A/C)* | *Chicken correlation (A/C)* | *Human A/C content (%)* | *Chicken A/C content (%)* |
|---|---|---|---|---|
| *ROBO1* | 0.381 | 0.402 | 20.1 | 21.5 |
| *PTPRK* | 0.437 | 0.407 | 23.2 | 22.2 |
| *FRY* | 0.396 | 0.323 | 22.5 | 19.1 |
| *CLEC16A* | 0.276 | 0.343 | 21.6 | 23.1 |
| *APBB2* | 0.395 | 0.477 | 23.1 | 22.9 |
| *ITSN1* | 0.401 | 0.469 | 23.6 | 20.0 |
| *AGAP1* | 0.391 | 0.34 | 22.3 | 20.5 |
| *EMC1* | 0.503 | 0.258 | 23.6 | 21.7 |
| *DOCK7* | 0.52 | 0.446 | 20.5 | 18.9 |
| *DCTN1* | 0.65 | 0.527 | 23.1 | 21.5 |
| *ASAP2* | 0.556 | 0.434 | 20.5 | 23.3 |
| *MEF2A* | 0.439 | 0.584 | 24.5 | 24.1 |
| *CPEB4* | 0.386 | 0.425 | 22.4 | 24.1 |
| *ROBO1* in cell | 0.455 | NA | 13.4 | NA |
| *EMC1* in cell | 0.194 | NA | 21.3 | NA |
| *DCTN1* in cell | 0.072 | NA | 27.8 | NA |
| *CPEB4* in cell | 0.239 | NA | 19.0 | NA |

**Supp Table S3. Cellular and *in vitro* structure probing correlations (Spearman).**

| *Gene* | *5NIA* | *DMS (A/C)* |
|---|---|---|
| *CPEB4* | 0.816 | 0.356 |

| | | |
|---|---|---|
| *EMC1* | 0.719 | 0.931 |
| *DCTN1* | 0.488 | 0.741 |
| *ROBO1* | 0.728 | 0.1 |

**Supp Table S4. Human and Chicken 5NIA correlation (Spearman).**

| *Gene* | *Correlation* |
|---|---|
| *ROBO1* | 0.47 |
| *PTPRK* | 0.272 |
| *FRY* | 0.269 |
| *CLEC16A* | 0.262 |
| *APBB2* | 0.327 |
| *ITSN1* | 0.278 |
| *AGAP1* | 0.392 |
| *EMC1* | 0.085 |
| *DOCK7* | 0.451 |
| *DCTN1* | -0.048 |
| *ASAP2* | 0.225 |
| *MEF2A* | 0.285 |
| *CPEB4* | 0.193 |

**Supplementary Table S5. Primers used for gene fragment amplification and cell probing.**

| *Name* | *Primer* |
|---|---|
| *T7_F* | TAATACGACTCACTATAGG |
| *Common_R* | TGTTGGAGTCACTCGACTCCGGT |
| *CPEB4_hsa_A_F* | CCCTACACGACGCTCTTCCGATCTNNNNNCAGTTGATTCTTTTGACAATGTTTGAAA |
| *CPEB4_hsa_A_R* | GACTGGAGTTCAGACGTGTGCTCTTCCGATCTNNNNNGGTCGTCTCCAAATACAAGTG |
| *CTN1_hsa_A_F* | CCCTACACGACGCTCTTCCGATCTNNNNNCCCGTAACCCCCAAATCACC |
| *DCTN1_hsa_A_R* | GACTGGAGTTCAGACGTGTGCTCTTCCGATCTNNNNNTTAGCTCCAACTCCCACCAGC |
| *EMC1_hsa_A_F* | CCCTACACGACGCTCTTCCGATCTNNNNNACTCTGCTAGGCAATGCACATTTT |
| *EMC1_hsa_A_R* | GACTGGAGTTCAGACGTGTGCTCTTCCGATCTNNNNNAGCCATGTGAGCAGGCAAC |
| *ROBO1_hsa_A_F* | CCCTACACGACGCTCTTCCGATCTNNNNNCTTTTCTGTTTCATTTTATGTTTCCTTATTT |
| *ROBO1_hsa_A_R* | GACTGGAGTTCAGACGTGTGCTCTTCCGATCTNNNNNCCAGTCACAGTGACACG |

### Supplemental Methods

#### Cell culture

Human medulloblastoma D341 Med (HTB-187) cells are from ATCC. Cells were maintained in Eagle's Minimum Essential Medium (EMEM; Gibco) supplemented with 20% fetal bovine serum (FBS) at 37°C with 5% CO2 and passaged once a week.

#### Cellular nascent RNA enrichment

~3 million D341 Med cells per sample collected with centrifugation at 500xg for 3 minutes. Cells were washed with 1ml cold supplemented 1X PBS (1ml 1X PBS, 1.33ul RNase inhibitor (NEB M0314L) and 1ul 1mM DTT added day of experiment). Cells were pelleted at 500xg for 5 minutes and resuspended in 750ul ice cold complete buffer A (incomplete: 10mM Tris-HCl, 300mM KCl, 10mM MgCl; 1.33ul RNase inhibitor and 1ul 1mM DTT added to 1ml incomplete buffer A day of experiment). 100ul of ice-cold complete lysis buffer (1% (v/v) Triton X-100, 0.2M sucrose, 1 Complete protease inhibitor cocktail tablet (Roche) per 30ml lysis buffer; 1.33ul RNase inhibitor and 1ul 1mM DTT added to 1ml incomplete lysis buffer day of experiment) was added and mixed with pipetting. Whole cell lysate was visualized with a countess after adding equal volume trypan blue to confirm sufficient cell lysis. Lysate was centrifuged at 500xg for 5 minutes at 4°C and the supernatant was discarded.

#### Cellular SHAPE probing

The nuclear pellet was resuspended in folding buffer (400 mM bicine pH 8, 200 mM NaCl, 20 mM MgCL2) and RNase inhibitor. The resuspended nuclear fraction was incubated with 25ul of 25 mM 5NIA or DMSO at 37°C for 5 minutes. RNA extraction was performed with TRIzol LS (Invitrogen, Thermo Fisher Scientific) according to the standard protocol. RNA was purified with the Monarch RNA Cleanup Kit (NEB) and underwent two rounds of DNase treatment (Turbo DNA Free Kit, Invitrogen) with double the amount of DNase and inactivation reagent described in the manufacturer's instructions. The RNA was cleaned up using RNA XP beads and 1ug was input into error-prone RT using SuperScript™ II with modified RT buffer (10X: 500 mM Tris pH 8, 750 mM KCl) supplemented with 6 mM manganese, RNAse inhibitor, random primers, DTT, and dNTPs. The RT product was cleaned up with RNA XP beads and split into 4 equal aliquots for amplicon library preparation. Two PCR reactions using gene specific primers were used to incorporate Illumina Truseq barcodes (Supp Table S5). Both PCR reactions included 15 cycles each with the Monarch Q5 High-Fidelity DNA Polymerase (NEB). Experiments were done in duplicate and paired-end libraries were sequenced at 2 x 150 on various Illumina platforms.

#### Cellular DMS probing

The nuclear pellet was resuspended in folding buffer (400 mM bicine pH 8, 200 mM NaCl, 20 mM MgCL2) and RNase inhibitor. The resuspended nuclear fraction was incubated with 25ul of 10% DMS or EtOH at 37°C for 5 minutes. 250ul 20% BME was added and incubated on ice for 5 minutes to inactivate the DMS. RNA extraction was performed with TRIzol LS according to the standard protocol. RNA purification, error-prone RT, and library prep was done the same way as with the Cellular SHAPE probing samples. Experiments were done in duplicate and paired-end libraries were sequenced at 2 x 150 on various Illumina platforms.

#### qRT-PCR for detection of chicken embryo sex

Quantitative reverse transcription polymerase chain reactions (qRT-PCRs) were performed on all RNA samples (post-DNAse treatment) to determine their sex so that equal numbers of both male and female embryos would be represented in our study. We used the iTaq Universal One-Step RT-qPCR Kit (Biorad) to perform these experiments and primer sequences were sourced from a recent publication which detailed a new method for using qRT-PCR on RNA isolated from chickens to determine biological sex (Subba et al. 2025). The primers used targeted either a sequence on the Z chromosome or two separate sequences on the W sex chromosome unique to heterogametic female birds (Subba et al. 2025). Sex was determined by empirical CT cutoff and confirmed by RNA sequencing analysis.
